# Predominance of positive epistasis among drug resistance-associated mutations in HIV-1 protease

**DOI:** 10.1101/822981

**Authors:** Tian-hao Zhang, Lei Dai, John P. Barton, Yushen Du, Yuxiang Tan, Wenwen Pang, Arup K. Chakraborty, James O. Lloyd-Smith, Ren Sun

**Affiliations:** Molecular Biology Institute, University of California, Los Angeles, CA 90095, USA; Shenzhen Institute of Synthetic Biology, Shenzhen Institutes of Advanced Technology, Chinese Academy of Sciences, Shenzhen 518055, People’s Republic of China; Department of Physics and Astronomy, University of California, Riverside, CA 92521, USA; School of Medicine, ZheJiang University, Hangzhou, 210000, China; Molecular and Medical Pharmacology, University of California, Los Angeles, CA 90095, USA; Department of Public Health Laboratory Science, West China School of Public Health, Sichuan University, Chengdu, China; Institute for Medical Engineering and Science, Departments of Chemical Engineering, Physics, & Chemistry, Massachusetts Institute of Technology, MA 21309, USA; Ragon Institute of MGH, MIT, & Harvard, Cambridge, MA 21309, USA; Department of Ecology and Evolutionary Biology, University of California, Los Angeles, CA 90095, USA; Fogarty International Center, National Institutes of Health, Bethesda MD 20892, USA

## Abstract

Drug-resistant mutations often have deleterious impacts on replication fitness, posing a fitness cost that can only be overcome by compensatory mutations. However, the role of fitness cost in the evolution of drug resistance has often been overlooked in clinical studies or *in vitro* selection experiments, as these observations only capture the outcome of drug selection. In this study, we systematically profile the fitness landscape of resistance-associated sites in HIV-1 protease using deep mutational scanning. We construct a mutant library covering combinations of mutations at 11 sites in HIV-1 protease, all of which are associated with resistance to protease inhibitors in clinic. Using deep sequencing, we quantify the fitness of thousands of HIV-1 protease mutants after multiple cycles of replication in human T cells. Although the majority of resistance-associated mutations have deleterious effects on viral replication, we find that epistasis among resistance-associated mutations is predominantly positive. Furthermore, our fitness data are consistent with genetic interactions inferred directly from HIV sequence data of patients. Fitness valleys formed by strong positive epistasis reduce the likelihood of reversal of drug resistance mutations. Overall, our results support the view that strong compensatory effects are involved in the emergence of clinically observed resistance mutations and provide insights to understanding fitness barriers in the evolution and reversion of drug resistance.

## 1 Introduction

Antibiotics and antiviral drugs have achieved great success in recent history [1]. However, therapeutic failure may occur due to low adherence and the emergence of drug resistance [2, 3]. The increasing amount of drug resistant pathogens is a global threat to public health [4–11]. The genetic barrier to drug resistance, defined as the number of mutations needed to acquire resistance, is a major determining factor of treatment outcomes [12–14]. Another important but often overlooked aspect of drug resistance is the fitness barrier [15–17]. Drug resistance associated mutations (DRAMs) in pathogen proteins may decrease enzymatic activities, interfere with molecular interactions, or destabilize the protein structure [18–22]. Because of the impaired replication capacity without drug selection, drug-resistant mutants cannot normally outcompete wild-type or establish in the population [23–25]. However, drug-resistant mutants can sometimes reach substantial frequency in the population. Fluctuating drug concentrations may create time windows when drug-resistant mutants replicate better than wild-type virus [26]. Moreover, compensatory mutations can rescue the impaired replication capacity of mutants and stabilize drug resistance [22, 27, 27–29]. Thus, comprehensive quantification of the fitness landscape is needed to predict the evolution of drug resistance [30, 31].

Epistasis, i.e. genetic interactions between mutations, is prevalent in molecular evolution [30–34]. Negative epistasis decreases fitness of the double mutant, posing constraints on gaining multiple mutations [35, 36]. It plays an important role in shaping the local fitness landscape [37]. Positive epistasis increases replication capacity of the double mutant, facilitating pathogens to acquire and maintain drug resistance [38–40]. Positive epistasis may create a fitness valley that prevents drug resistant mutations from reversal [41]. Collectively, positive and negative epistasis determine the topography of the fitness landscape [42] and the course of drug resistance evolution [32]. Empirical studies on the genetic interactions between DRAMs, especially in high-order mutants, are still rare [43, 44].

HIV-1 protease inhibitors are important components of combination antiretro-viral therapy [45] that target HIV-1 protease enzymatic activity [46, 47]. Second-generation protease inhibitors have extremely high binding affinity to viral protein [48]. Resistance to them typically requires more mutations than resistance to first-generation protease inhibitors and other antiretroviral drugs [49,50]. For example, mutation K103N on reverse transcriptase is sufficient to confer HIV-1 nevirapine (NVP) resistance [51], while more than 4 *de novo* mutations are needed for protease inhibitor darunavir (DRV) resistance [52]. Protease inhibitor-resistant viruses with multiple DRAMs also have significantly reduced fitness [53, 54]. HIV-1 gained DRAMs on protease during sub-optimal protease inhibitor therapy [55]. Most resistance mutations directly affect the binding affinity between HIV-1 protease and the inhibitor, but they are likely to be deleterious because they also reduce binding to the native substrate of HIV-1 protease. To compensate the deleterious effect, some other DRAMs stabilize HIV-1 protease, allowing drug-resistant virus to replicate as efficiently as its parental wild-type virus [27, 56]. Meanwhile, reversals of protease inhibitor resistance-associated mutations were rarely seen clinically, even when therapy was interrupted [57] or when mutant virus infected drug-naïve patients [58, 59]. These observations indicate that epistasis may be important for the evolution of protease inhibitor resistance.

Here, we present a quantitative high-throughput genetics approach [60, 61] to study the fitness distribution and epistasis of HIV-1 protease inhibitor DRAMs. Combining these data with clinical data and fitness models, we found that positive epistasis was predominant and especially enriched among DRAMs, and prevalent along drug resistance evolutionary paths. Our results suggest that fitness hills created by epistasis result in barriers that entrench DRAMs, and thus drug-resistant viruses are unlikely to revert after transmission to drug-naïve patients or discontinuation of anti-retroviral drug treatment.

## 2 Results

### 2.1 Fitness profiling of DRAMs in HIV protease

To study the interactions among DRAMs in HIV protease, we constructed a library of virus mutants that covers combinations of amino acid substitutions at 11 resistance-associated sites in HIV protease (Figure 1A, Table 1, 2^9^ × 3^2^ = 4608 genotypes). These sites have been annotated as major drug resistance sites in Stanford Drug Resistance Database [62,63], and all have been shown to be strongly associated with drug resistance [3]. In our mutant library, 9 sites have one amino acid substitution and the other 2 sites have 2 amino acid substitutions (Figure 1A, Table 1). 2736 out of 4608 possible genotypes (59.38%) were covered in the plasmid library.

**Table 1:**
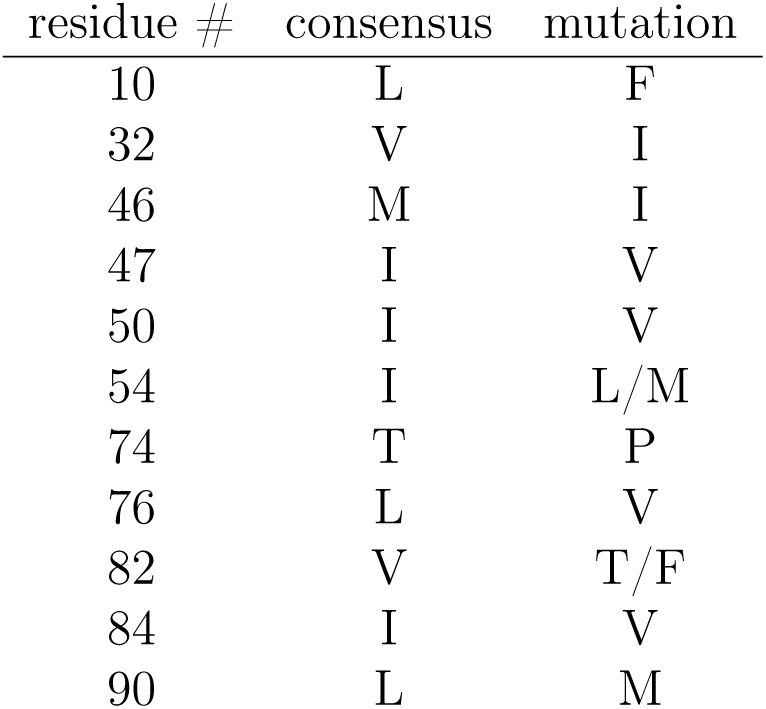
List of protease inhibitor resistance associated mutations covered in the library.

| residue # | consensus | mutation |
| --- | --- | --- |
| 10 | L | F |
| 32 | V | I |
| 46 | M | I |
| 47 | I | V |
| 50 | I | V |
| 54 | I | L/M |
| 74 | T | P |
| 76 | L | V |
| 82 | V | T/F |
| 84 | I | V |
| 90 | L | M |

**Figure 1.**
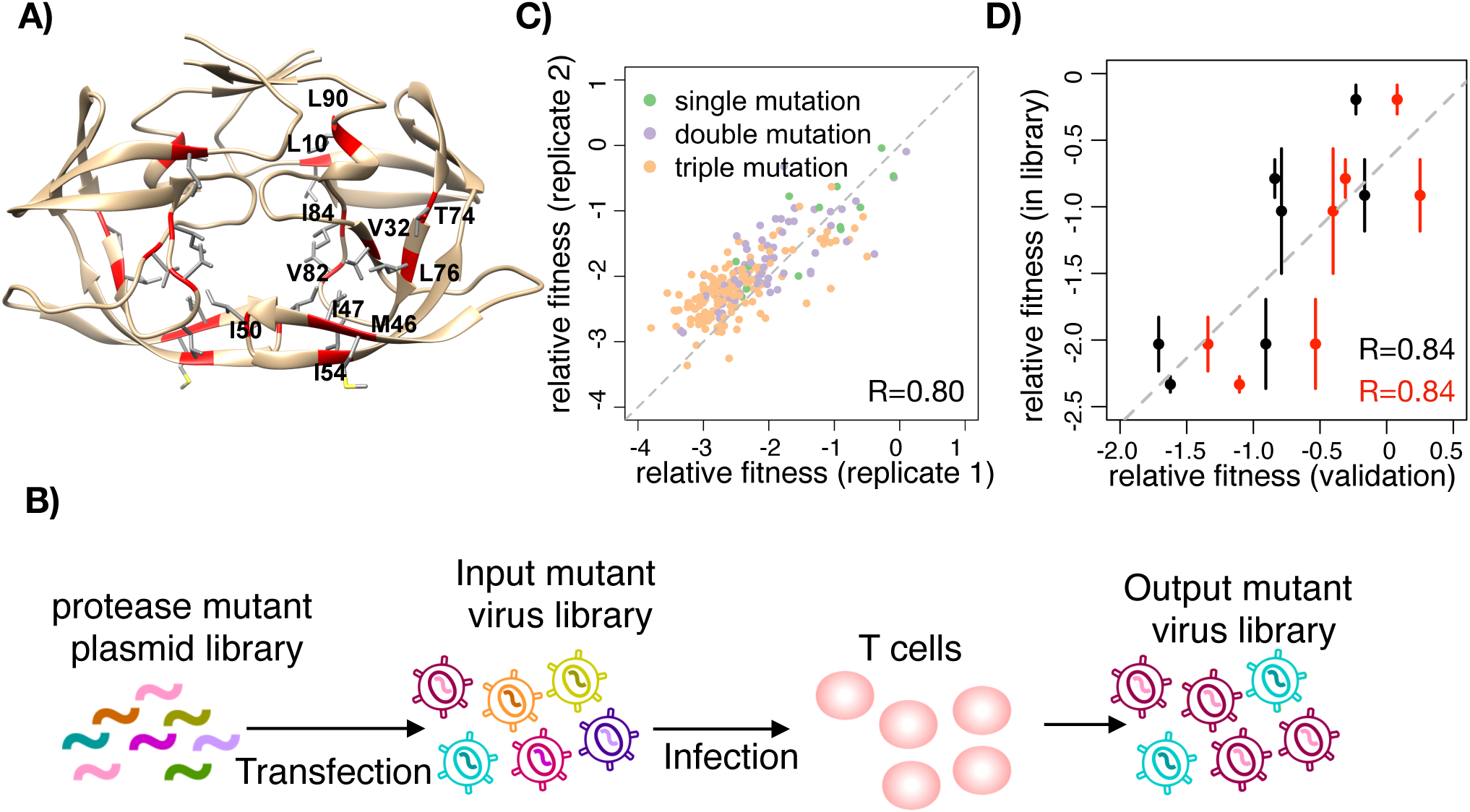
High-throughput fitness profiling of combinatorial HIV-1 protease mutant library. A) The structure of protease dimer (PDB: 4LL3). The side chains of drug resistance associated residues are shown. B) Protease mutations were introduced into NL4-3 background. T cells were infected by the mutant virus library. The frequency of mutants before (input library) and after (output library) selection were deep sequenced. C) The correlation of relative fitness between two biological replicates. Pearson correlation coefficient (R) is 0.80. D) Two independent validation experiments were performed. We constructed 7 protease single mutant plasmids and recovered viruses independently. We mixed each mutant virus with wild-type virus (validation 1, black dots) and passaged in T cells for 6 days. We also mixed all 7 mutant viruses together with wild-type (validation 2, red dots) and infected T cells for 6 days. The relative fitness of each mutant was quantified by the same means as that in the library. Pearson correlation coefficients (R) for validation 1 and validation 2 are both 0.84. Error bar is standard deviation (n=3).

We quantified the relative fitness of mutants using high-throughput fitness profiling (Figure 1B, See Methods for details). We performed 3 independent transfection experiments to validate the reproducibility of fitness profiling. For each biological replicate, relative fitness was calculated independently. The Pearson’s correlation coefficients of single, double and triple mutations between replicates range from 0.80 to 0.82 (Figure 1C and Figure S1). After filtering out mutants with low frequency or low reproducibility among replicates of input virus libraries (see Methods for details), we were able to estimate the relative fitness of 1219 genotypes. The fitnesses of all single mutants, and more than 70% of double and triple mutants, were quantified (Figure S2).

To validate the quantification of relative fitness, we conducted competition experiments with individually constructed protease mutants. We performed two sets of validation experiments. For the first set, we packaged mutant virus and wild-type virus independently and mixed them in pairs for head-to-head competition. The frequency of mutant virus and wild-type virus were quantified by deep sequencing and the relative fitness was calculated in the same way as we did in library screening. A total of 7 mutants were constructed and validated. For the second set of experiments, we mixed all 7 single mutants with wild-type virus in competition experiments. The relative fitness was defined in the same way. The fitness measured in validation experiments was highly correlated with the fitness in library screening (Figure 1D, *R* = 0.84 for each independent validation, Pearson’s correlation test).

### 2.2 Positive epistasis rescues the mutational load of DRAMs

We first looked at fitness effect of DRAMs. In our definition, a mutant virus of relative fitness −1 means that the relative frequency of this mutant drops 10 fold after infection in cell culture. All single mutations were deleterious to virus replication (Figure 2A). The relative fitness of single mutants ranged from −2.33 (V82F) to −0.19 (L90M). This is consistent with previous reports that randomly introduced mutations were mostly deleterious to protease enzymatic activity or HIV-1 replication capacity [34, 64–66]. Random mutagenesis in other viruses also revealed a lack of beneficial mutations in well-adapted systems [65, 67–69]. DRAMs in particular were also reported to be deleterious to virus replication [31, 44]. They may destabilize viral protein, affect enzymatic activities or impact other protein-protein interactions [21, 70].

**Figure 2.**
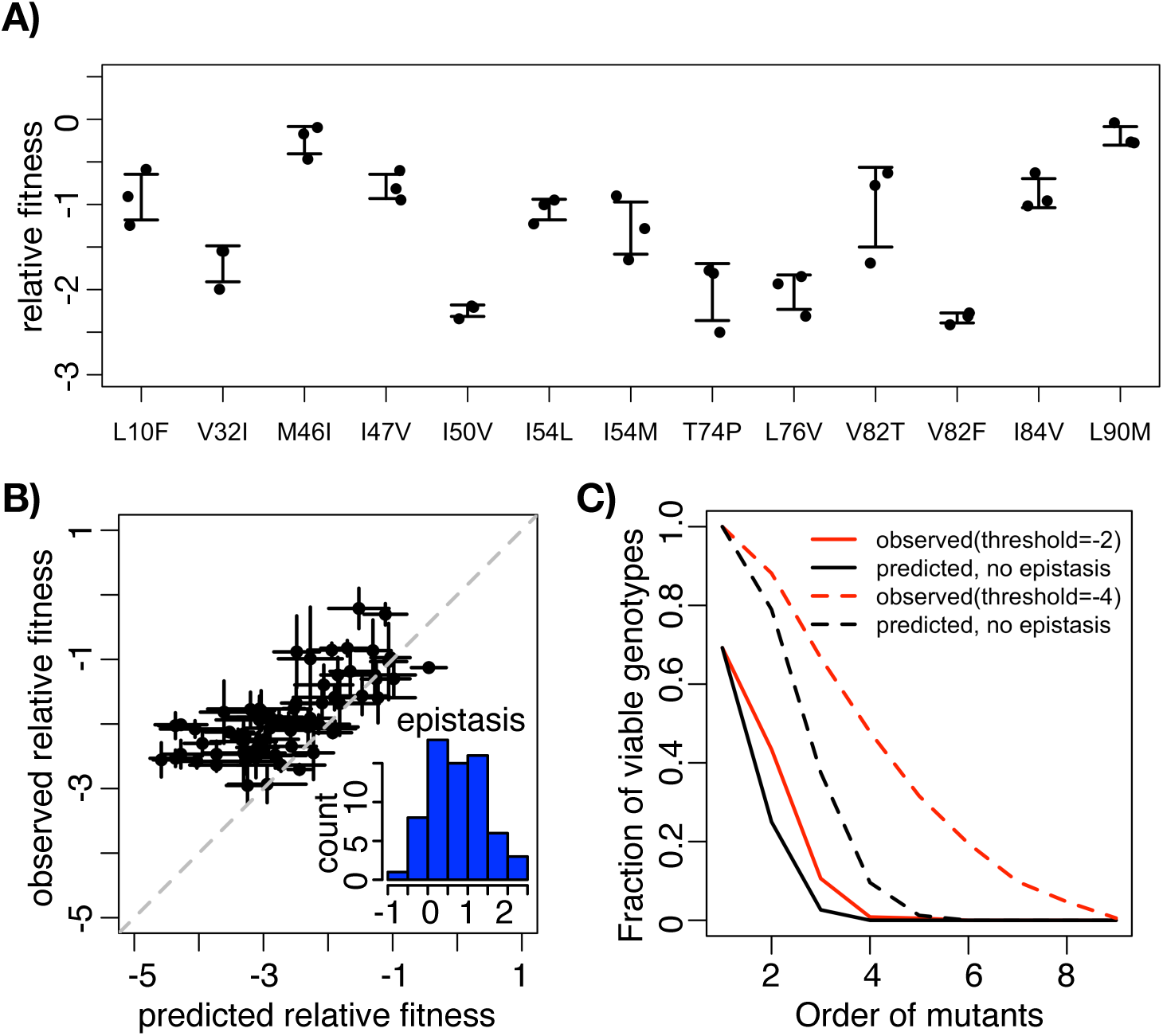
Positive epistasis rescues the mutational load of DRAMs. A) Relative fitness of single mutants. Error bar is standard deviation (n=3). B) The predicted relative fitness and observed relative fitness of double mutants. The predicted relative fitness was the sum of that of the two single mutants. Inset, the distribution of epistasis between double mutants. Error bar is standard deviation (n=3). C)The predicted and observed fraction of viable mutants. A mutant was defined as viable if its relative fitness is higher than −4(dashed line) or −2(solid line).

We then analyzed epistasis between all pairs of DRAMs. Previous studies have shown the prevalence of epistasis among pairs of random mutations [34, 37, 67] or spontaneously accumulated mutations [71]. However, studies focused on the epistasis among drug resistance mutations are still limited [30, 39, 64, 67, 72]. Based on the fitness effect of single DRAMs, we predicted the relative fitness of double mutants with the assumption that no epistasis existed among any two single mutations (i.e., the predicted relative fitness of a double mutant was the sum of those of two single mutants)(Figure 2B). Surprisingly, the observed relative fitness of most double mutants were significantly higher than the predicted values (*p* = 2.2 × 10^−6^, twosided Wilcoxon rank sum test), suggesting that positive epistasis is prevalent among DRAMs (Figure 2B inset). Pairwise epistasis between two DRAMs is quantified as *ε*_*i,j*_ = *f*_*i,j*_ −*f*_*i*_ −*f*_*j*_, *f*_*i*_ represents the relative fitness of mutants *i*. The distribution of epistasis ranged from −0.69 (M46I and L90M) to 2.34 (L76V and V82F) and 86.6% of pairwise interactions between DRAMs are positive.

We also analyzed the extent of epistasis among high-order mutants. We observed a trend that relative fitness decreased as the order of mutants increased (Figure S3). This is consistent with previous reports that mutational load restricted virus replication capacity [30,73,74]. To better quantify the fitness cost of multiple mutations, we calculated the frequency of viable mutants by different thresholds, *f* > −2 or *f* > −4. The frequency of viable mutant virus decreased as the number of mutations increased (Figure 2C), consistent with previous observations in HIV-1 and other RNA viruses [75–78]. We then predicted the relative fitness of high-order mutants by summing the relative fitness of corresponding single mutants. We observed more viable mutants than would be predicted without epistasis (Figure 2C).

This indicated pervasive positive epistasis rescued high-order mutants from lethal relative fitness, which is consistent with other clinical observations in protease inhibitor resistant virus [30,44]. As a result, positive epistasis partially relieved HIV-1 mutational load and allowed viruses to explore more sequence space.

### 2.3 Enrichment of positive epistasis among DRAMs

There are two possible explanations for the observed positive epistasis among DRAMs of HIV protease. The first hypothesis is that all mutations in HIV protease tend to interact positively. The second hypothesis is that epistasis among random mutations in HIV protease is on average zero, but positive epistasis is enriched among DRAMs. We introduced the Potts model to test our hypotheses, while simultaneously testing whether our finding of prevalent positive epistasis among DRAMs carries over to the clinical setting. Potts models, originally developed in statistical physics, have been employed previously to use the population-level frequencies and correlations between different mutations to estimate their fitness effects [79–82]. In the Potts model, the probability of observing a genotype 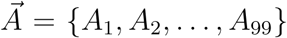 is given by equations in Figure 3A. Here the *A*_*i*_, *i* ∈ {1, 2,…, 99} are variables that represent the amino acid at site *i* on each of the 99 sites of protease. Two sets of Potts parameters, fields *h*_*i*_(*A*_*i*_) and couplings *J*_*ij*_(*A*_*i*_, *A*_*j*_), give the statistical energy 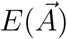, which is negatively correlated with fitness. These parameters are estimated in order to reproduce the frequencies and correlations between mutations that are observed in the data. The fields *h*_*i*_(*A*_*i*_) represent the fitness effect of amino acids *A*_*i*_ at sites *i* alone, while the couplings *J*_*ij*_(*A*_*i*_, *A*_*j*_) describe epistatic interactions between amino acids *A*_*i*_ at site *i* and *A*_*j*_ at site *j*. For both the couplings and the fields, positive parameter values correspond to beneficial effects on fitness, while negative values correspond to deleterious fitness effects. We applied a maximum entropy method [83] to an alignment of 20911 HIV-1 clade B protease sequences from drug naïve patients, obtained from the Los Alamos National Laboratory HIV sequence database (hiv.lanl.gov, accessed 24 March 2017) to calculate these two sets of Potts parameters.

**Figure 3.**
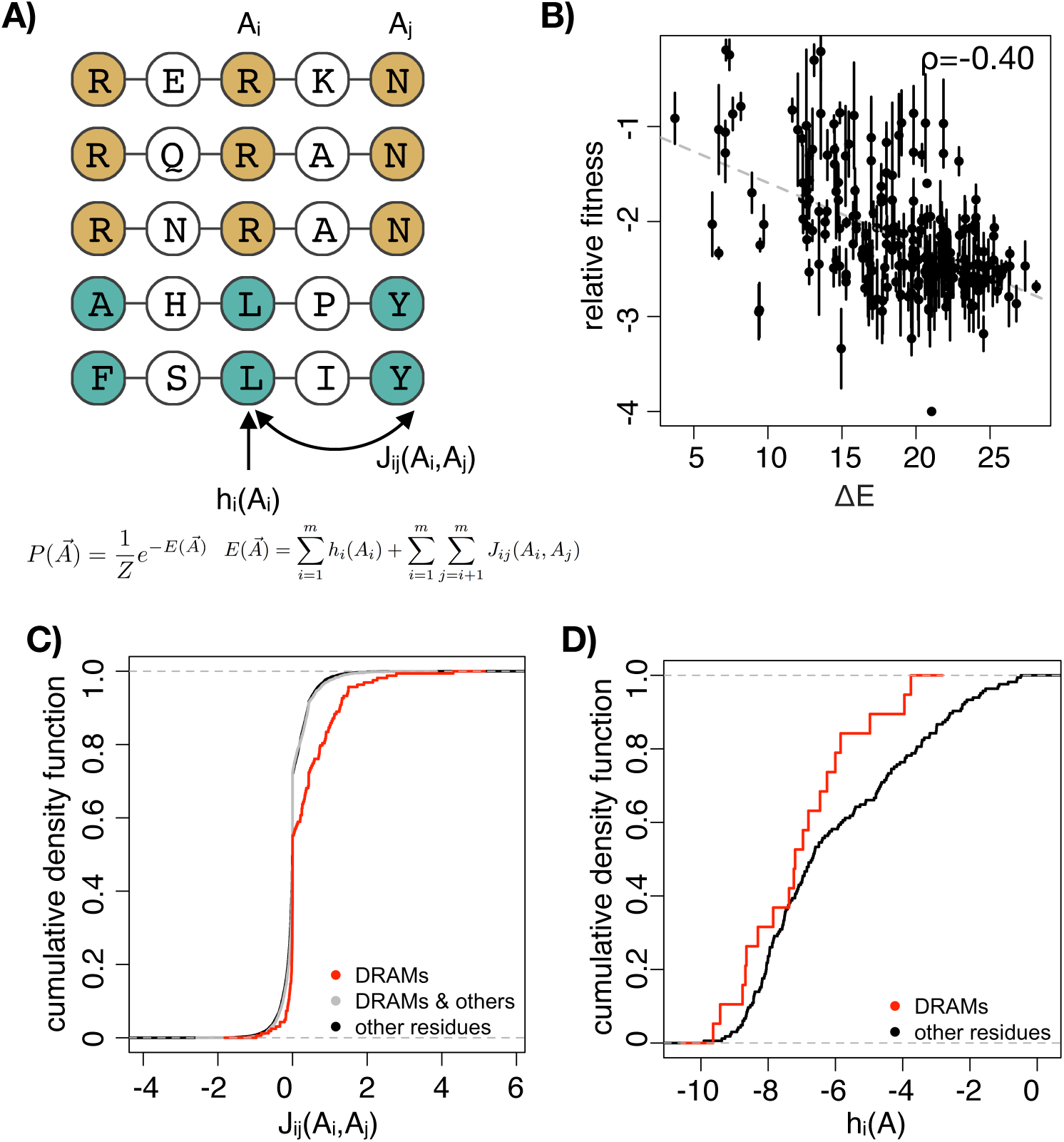
Positive epistasis is enriched among DRAMs. A) The conceptual graph of Potts model. Potts model uses the probability of mutations occurring with other mutations to estimate the statistical energy. h_i_ is the field parameter while J_ij_ is the coupling parameter. B) The correlation of Potts energy(ΔE = E_mut_-E_WT_) and relative fitness of mutations with lower than 4 DRAMs. Spearman correlation coefficient (ρ) is −0.40. C)The cumulative density function of coupling parameters of DRAMs and all other mutations. Coupling parameters between DRAMs are more positive positive than those between DRAMs & others (D = 0.22, p = 2.1×10^−7^, two-sided K-S test) and those between other residues(D = 0.22, p = 5.1×10^−7^, two-sided K-S test). D) The cumulative density function of field parameters of DRAMs and all other mutations. Field parameters of DRAMs and other residues are not significantly different(D = 0.25, p = 0.20, two-sided K-S test).

Then we calculated 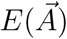 for all mutants in our protease library. We found that the Potts energy (Δ*E* = *E*_*mut*_ − *E*_*WT*_) is significantly correlated with the relative fitness we measured in our screening (*ρ* = −0.40, *p <* 2.2 × 10^−16^, Spearman’s correlation test, Figure 3B). The correlation was lower than previous analysis in HIV-1 Gag and Env region [80, 82]. This may be due in part to strong phylogenetic bias on the inferred Potts parameters, because protease is particularly highly conserved. It is also possible that epistatic interactions with other parts of the HIV-1 genome and complicated anti-innate immunity functions of protease [84] obscure the effects of individual mutations on replicative fitness in vitro.

We compared the couplings between DRAMs and other mutations in protease. The Potts couplings *J*_*ij*_(*A*_*i*_, *A*_*j*_) give the contribution of pairwise epistatic interactions between amino acids *A*_*i*_ and *A*_*j*_ at sites *i* and *j*, respectively. We compared the couplings among DRAMs and among all other possible mutations on protease (Figure 3C). Couplings of other protease mutations clustered near 0, while those of DRAMs are significantly more positive than that of other mutations (D = 0.22, *p* = 2.1 × 10^−07^, two-sided K-S test). Moreover, *J*_*ij*_(*A*_*i*_, *A*_*j*_) among DRAMs were also more positive than those between DRAMs and other residues (D = 0.22, *p* = 5.1 × 10^−07^, two-sided K-S test). Although the fields *h*_*i*_(*A*_*i*_) of DRAMs are more negative than other mutations, the difference is not significant (Figure 3D, D = 0.25, *p* = 0.20, two-sided K-S test). Overall, analysis based on the Potts model is consistent with our experimental results that positive epistasis is enriched among DRAMs, and lends support to our second hypothesis that epistasis among random mutations in HIV protease is on average zero.

### 2.4 Implications of positive epistasis in evolution

To study the role of epistasis in evolution, we analyzed the evolutionary pathways covering all genotypes with up to 4 amino acid substitutions from the wild-type virus (13 single mutants, 67 double mutants, 176 triple mutants and 290 quadruple mutants) (Figure 4A). Mutants are linked if they differ by one amino acid substitution. We have found that all 13 DRAMs are deleterious on the wild-type background (Figure 2A). However, the fitness effect of a single DRAM becomes less deleterious on genetic backgrounds where other DRAMs have been fixed (Figure S4). Following the generalized definition of epistasis proposed by Shah et al. [85], we define trajectory-based epistasis *ε*_*M,j*_ that measures the deviation of the fitness effect if the order of mutations were reversed. *ε*_*M,j*_ = *f*_*M,j*_ − *f*_*M*_ − *f*_*j*_, where *f*_*M*_ and *f*_*j*_ represent the relative fitness of background *M* and single mutant *j* [86]. For example, mutation *j* can be deleterious on the wild-type background but beneficial on another genetic background that mutation *i* has been fixed. Trajectory-based epistasis is calculated for each amino acid substitution and averaged over genetic backgrounds with a certain Hamming distance to the wild-type (Figure 4B). For all DRAMs profiled in this study, we find that trajectory-based epistasis is overall positive and increases steadily with the number of substitutions, i.e. the fitness contribution of a single mutations becomes more positive if more DRAMs have been fixed.

**Figure 4.**
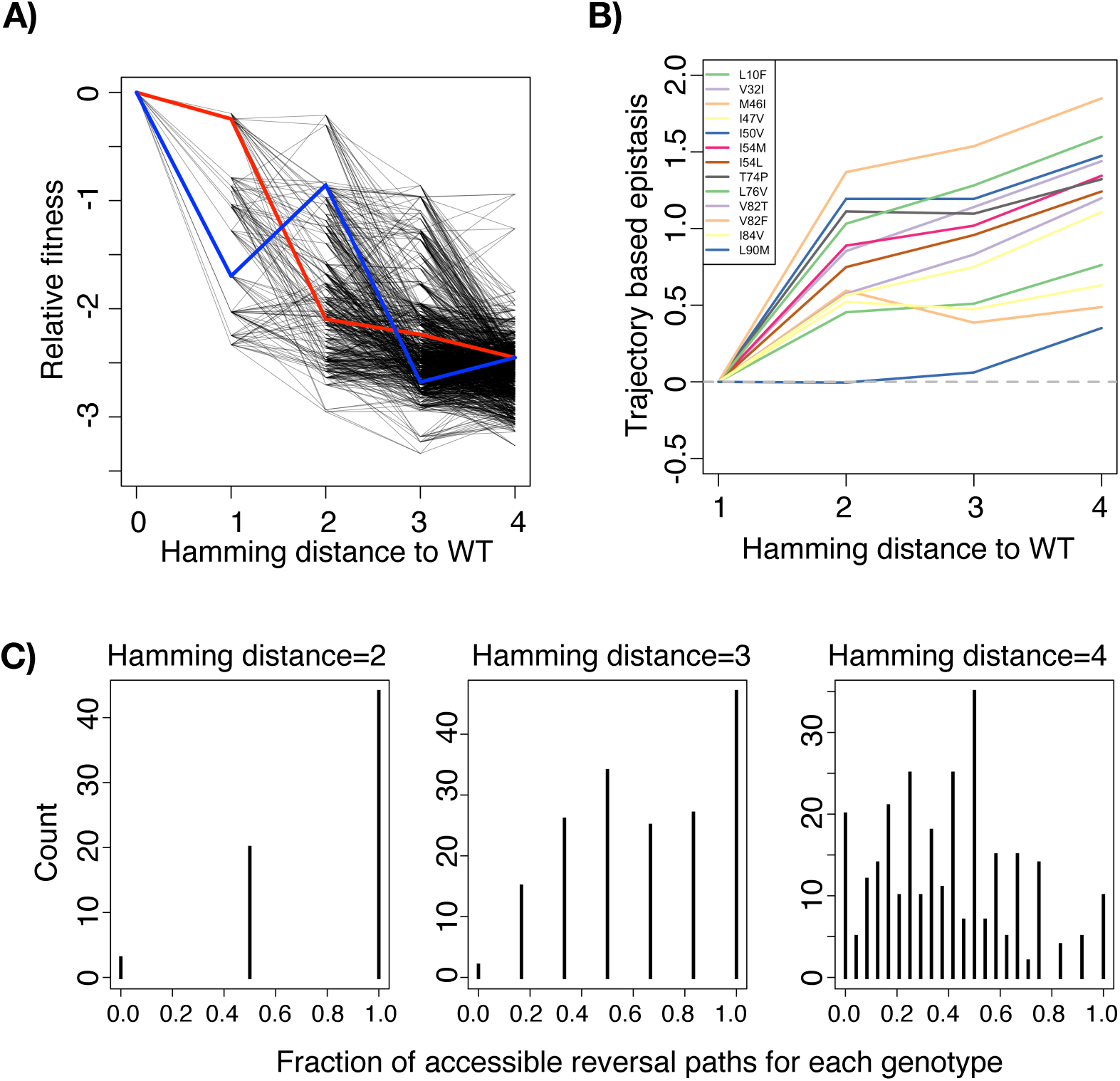
Ruggedness in fitness landscapes prevents DRAMs from reversion to wild-type. A) Fitness with possible evolutionary trajectories. Mutants are linked if they only have one residue difference. Red line represents an accessible path that a quadruple mutant can take and reverse to wild-type. Blue line represents an inaccessible reversal path to wild-type for that mutant. B) Trajectory-based epistasis is calculated for each amino acid substitution and averaged over genetic backgrounds with a certain Hamming distance to the wild-type. The fitness effect of a single mutation becomes less deleterious on genetic backgrounds where other DRAMs have been fixed. C) The distribution of accessible paths for all genotypes with a certain hamming distance to wild type.

We tested the hypothesis that positive epistasis prevented drug resistance associated genotypes from reverting to wild-type [41,87,88]. Although DRAMs incurred significant fitness cost, some drug resistant mutants would not revert to wild-type after transmitting to a drug naïve patient. We quantified the frequency of accessible evolutionary pathways between mutants and wild-type in our experimentally measured fitness landscape of HIV protease DRAMs. A reversal path is defined to be accessible if and only if the virus fitness increases monotonically along the path. For example, quadruple mutant V32I_M46I_I54L_V82F has many paths to revert to wild-type (Figure 4A). Among them, losing V32I, I54L, V82F, M46I in order was an accessible path (Figure 4A, red line). On the contrary, losing I54L, V82F, M46I, V32I is not accessible because there were 2 steps with decreasing relative fitness (Figure 4A, blue line). We found that among double mutants, 44 have two accessible reversal paths to the wild type, 20 have only one accessible reversal path, and interestingly 3 of them have none. These 3 mutants (I50V_T74P, M46I_I54M and L76V_V82F) represent local fitness peaks and the reversal to wild-type is blocked by a fitness valley. We found that the number of accessible reversal paths decreased with the accumulation of DRAMs (Figure 4C). This indicates that protease mutants become less likely to revert to wild-type as the number of DRAMs increases. Our results are consistent with clinical observations that protease inhibitor resistance associated mutations seldom reverted even when therapies were interrupted [25, 57] or drug-naïve patients were infected [58, 59]. The difficulty of reversal also explains the rising frequency of drug resistant HIV-1 viruses in acute phase patients [41, 88].

## 3 Discussion

In this study, we systematically quantified the fitness effect of DRAMs of HIV-1 protease. While all DRAMs reduced the virus replication fitness, pervasive positive epistasis among DRAMs alleviated the fitness cost substantially. Moreover, we analyzed the HIV sequence data from patients by the Potts model. We found the statistical energy inferred from HIV sequences *in vivo* correlated well with the replication fitness measured *in vitro*. Based on our fitness data and the mutational couplings inferred by the Potts model, we showed that positive epistasis is enriched among DRAMs of HIV-1 protease, in both local fitness landscape and evolutionary paths. Finally, we studied the role of epistasis in evolutionary pathways. We found that positive epistasis among DRAMs entrenches drug resistance and blocks the reversal paths to wild-type virus, which has important implications for the design of anti-retroviral therapies.

There are a few limitations of this study. Firstly, we only measured the fitness effect of DRAMs in the absence of protease inhibitors. We are not able to quantify drug resistance of DRAMs because protease inhibitors block multiple rounds of virus infection and prevent us from accurate examination of mutant frequency under drug selection. Also, we did not sequence other genes of HIV-1. HIV-1 mutates rapidly due to low fidelity of reverse transcriptase [89, 90]. There might be compensatory mutations occurring on other proteins that rescued the protease DRAMs. Thirdly, the correlation between our validation experiments and high-throughput screening experiments was less than correlations from bacteria and yeast experiments [91, 92]. Mechanistic difference between logistic growth and viral growth may complicate the quantification of viral fitness [93]. Direct measurement of viral frequency may not linearly correlate to the probability of replication [94].

Statistical models suggest a pervasive negative distribution of fitness effect for single mutations on HIV-1 [31, 80, 95]. Previous models also predicted the entrenchment of deleterious DRAMs by positive epistasis [96, 97]. This dataset provides a unique chance to experimentally test these statistical hypotheses. The predominance of positive epistasis is also observed in HIV-1 [30] and in other organisms [39, 98]. However, they either relied on naturally-occurring resistant clones or indirectly activating gene functions. This report is the first dataset to systematically quantify the epistasis among functional residues in HIV-1 drug resistance evolution, without the bias of drug selection and *in vivo* evolution. Overall, our results are important for understanding drug resistance evolution. We found positive epistasis plays a critical role in HIV-1 gaining and maintaining drug resistance. Epistasis makes the fitness landscape rugged, preventing DRAMs from reversion to wild-type, even when antiviral therapy is interrupted or virus transmits to a healthy individual [87, 99].

Positive epistasis involves many kinds of molecular mechanisms. We find that the relative fitness of single mutants is not a significant factor of positive epistasis. We compared *h*_*i*_ in the Potts model for all DRAMs and other single mutants. They were not significantly different(*p* = 0.20, K-S test). Physical distance between residues is a significant factor contributing to positive epistasis. The physical distances between these residues were significantly less than those between any two random residues on HIV-1 protease (D = 0.32, *p* = 3.9 × 10^−10^, two-sided K-S test, Figure S5A), suggesting that physical contact among DRAMs might contribute to the observed positive epistasis. Some mutations may have structurally stabilizing effect to other residues. We used FoldX to predict the folding free energy (ΔΔ*G*) as a quantification of protein stability [100] for all mutants in our library (Figure S5B). We noticed in the screening that mutation V82F contributed to the positive epistasis on many genetic backgrounds (Figure 4B), but it did not contribute much to the stabilizing effect. Thus, structurally stabilizing effects did not completely explain the positive epistasis among these mutations. Other mechanisms like protease inhibitor binding affinity and substrate specificity may also contribute to the epistasis among these mutations.

## 4 Material and Methods

### 4.1 Plasmid library construction

HIV-1 DRAMs were picked according to their prevalence in protease inhibitor treated patients [3]. We chose 11 residues with 13 mutations to construct a combination of HIV-1 protease mutant library (Table 1).

We used a ligation-PCR method to construct the library on NL4-3 backbone, which is an infectious subtype B strain. All possible combinations of these 13 mutations are 2^9^ × 3^2^ = 4608 genotypes. The mutagenesis region spanned 243 nucleotides on HIV-1 genome. We split the region into 5 oligonucleotides and ligate them in order by T4 ligase (from New England BioLabs). The sequence of oligonucleotides are shown in Supplementary File 2. After each ligation, we recovered the product by PCR and used restriction enzyme BsaI-HF (from New England BioLabs) to generate a sticky end for the next step ligation.

After making the 243-nucleotide mutagenesis fragment, we PCR amplified the upstream and downstream regions near this fragment and used overlap extension PCR to ligate them together. We then cloned it into full length HIV-1 NL4-3 background. We harvested more than 30,000 *E. coli* colonies to ensure sufficient coverage of the library complexity.

### 4.2 Virus production

The plasmid DNA was purified by HiPure Plasmid Midi Prep Kit (from Thermo Fisher Scientific). To produce virus, we used 16 µg plasmid DNA and 40 µL lipofectamine 2000 (from Thermo Fisher Scientific) to transfect 2 × 10^7^ 293T cells, in 3 independent biological replicates. We changed media 12 hours post transfection. The supernatant was harvested 48 hours post transfection, labeled as input virus and frozen at −80 °C. Virus was quantified by p24 antigen ELISA kit (from PerkinElmer).

### 4.3 Library screening

CEM cells were cultured in RMPI 1640 (from Corning) with 10% FBS (from Corning). To passage library in T cells, we added 25 mL viruses and 120 µg polybrene to 50 million CEM cells. We achieved 10 ng p24 for every million CEM cells. We washed cells and completely changed media 6 hours post infection. We supplemented the cells with fresh media 3 days post infection and harvested supernatant 6 days post infection. We centrifuged supernatant at 500 × *g* for 3 minutes to remove the cells and cell debris. The rest of supernatant was frozen at −80 °C.

### 4.4 Sequencing library preparation

We used QIAamp viral RNA mini kit (from QIAGEN) to extract virus RNA from supernatant. We then used DNase I (from Thermo Fisher Scientific) to remove the residual DNA. We used random hexamer and SuperScript III (from Thermo Fisher Scientific) to synthesize cDNA. The virus genome copy number was quantified by qPCR. The qPCR primers are 5’-CCTTGTTGGTCCAAAATGCGAAC-3’ and 5’ATGGCCGGGTCCCCCCACTCCCT-3’.

At least 2 × 10^5^ copies of viral genome were used to make sequencing libraries. We PCR amplified the mutagenesis regions using the following primers: 5’-CTAA TCCTGGAGTCTTTGGCAGCGACCC-3’ and 5’-GAAGACCTGGAGTGCAGCCAATCTGAGT-3’. We then used BpmI (from New England BioLabs) to cleave the primers and ligate the sequencing adapter to the amplicon. We used PE250 program on Illumina MiSeq platform to sequence the amplicon.

### 4.5 Calculation of fitness and epistasis

We used custom python codes to map the sequencing reads to reference NL4-3 genome. Mutations were called if both forward and reverse reads have the same mutation and phred quality scores are both above 30. All codes are available on https://github.com/Tian-hao/protease-inhibitor. All data were deposited in SRA (short read archive) database under accession PRJNA546460. For each replicate of the virus library from the transfected 293T cells, we reached 4.45 × 10^5^ to 6.05 × 10^5^ sequencing depth. We filtered out the genotypes with frequency fewer than 5 × 10^−5^ in any biological replicate and the genotypes whose frequency differ more than 10 folds between any two biological replicates.

Relative fitness *f*_*m,r*_ of mutant *m* in experiment *r* (biological replicates) was defined as Equation 1.

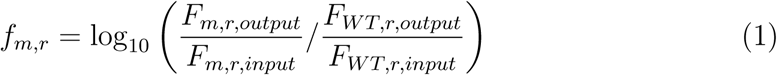

*F*_*m,r,input*_ is the frequency of mutant *m* before screening. *F*_*m,r,output*_ is the frequency of mutant *m* after passaging. *F*_*WT,r,input*_ is the frequency of wild-type virus before screening. *F*_*WT,r,output*_ is the frequency of wild-type virus after passaging.

The relative fitness *f*_*m*_ was defined as the average of 3 biological replicates (Equation 2). However, if relative fitness was missing in one replicate, we only average the other two replicates. The relative fitness value of all mutants was shown in Supplementary File 1.

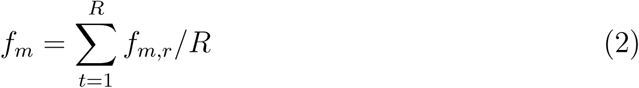

where *R* is the number of biological replicates.

Pairwise epistasis *ε*_*i,j*_ between mutant *i* and mutant *j* was defined as:

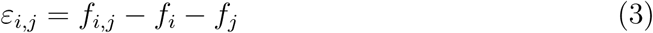

where *f*_*i,j*_ refers to the relative fitness of double mutant *i* and *j*.

Trajectory-based epistasis *ε*_*M,j*_ between a multi-mutation genotype *M* and another genotype differ by one mutation *j* was defined as:

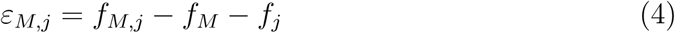

### 4.6 Potts model

Data used to infer parameters for the Potts model were downloaded from the Los Alamos National Laboratory HIV sequence database, as described in the main text. Sequences were processed as previously described [101]. Briefly, we first removed insertions relative to the HXB2 reference sequence. We also excluded sequences labeled as “problematic” in the database, and sequences with gaps or ambiguous amino acids present at >5% of residues were removed. Remaining ambiguous amino acids were imputed using simple mean imputation.

Each sequence in the multiple sequence alignment (MSA) is represented as a vector of variables 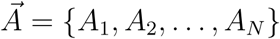, where *N* = 99 is the length of the sequence. Each of the *A*_*i*_ represents a (set of) amino acid(s) present at residue *i* in the protein sequence. To choose the amino acids at each site that would be explicitly represented in the model, we first computed the frequency 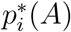 of each amino acid *A* at each site *i* in the MSA. To compute these frequencies, we weighted the sequences such that the weight of all sequences from each unique patient was equal to one, thereby avoiding overcounting in cases where many sequences were isolated from a single individual. We then explicitly modeled the *q*_*i*_ most frequently observed amino acids at each site that collectively capture at least 90% of the Shannon entropy of the distribution of amino acids at that site [101]. All remaining, rarely observed amino acids were grouped together into a single aggregate state. For these data, this choice resulted in an average of three explicitly modeled states at each site (minimum of 2, maximum of 6).

The Potts model is a probabilistic model for the ‘compressed’ sequences 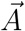, where the probability of observing a sequence 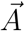 is

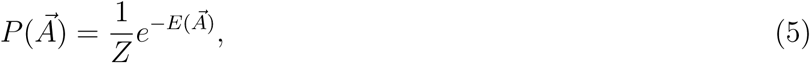

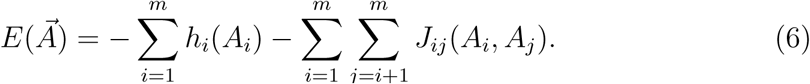

Here the normalizing factor

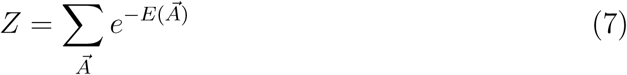

ensures that the probability distribution is normalized. We used ACE [83] to infer the set of Potts fields *h*_*i*_(*A*_*i*_) and couplings *J*_*ij*_(*A*_*i*_, *A*_*j*_) that result in average frequencies and correlations between amino acids in the model (5) that match the frequencies 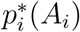 and correlations 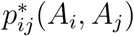 observed in the data. We used a regularization strength of *γ* = 7 × 10^−5^ in the inference, which is roughly equal to one divided by the number of unique patients from which the sequence data were obtained. We confirmed that the parameters inferred by ACE resulted in a Potts model that accurately recovered the correlations present in the data.

### 4.7 Validation experiments

We constructed 7 single mutants by site-directed mutagenesis. The primers used this experiment are listed in Supplementary File 2. We used overlap-extension PCR to amplify the fragment with mutated nucleotides. We ligated the fragment with NL4-3 backbone using ApaI and SbfI. We transformed competent *E*.*coli* and picked single colonies. We sequenced the protease region of plasmids to make sure there is only desired mutant in this region. 7 mutants were L10F, I47V, T74P, L76V, V82F, V82T, L90M.

We produced mutant viruses in 293T cells, mixed them with wild-type and infected CEM cells. The frequencies of mutant virus before and after infection were quantified by deep sequencing. We did 2 biological replicates with each validation method. For validation 1, we pairwisely mixed the mutant and wild-type virus for competition. For validation 2, we mixed all 7 mutants and wild-type virus.

## Ethics Statement

Reagents were acquired from the NIH AIDS Reagent program. The work is approved by UCLA IRB.

## 5 Author contributions

L.D., R.S., J.L.S. and T.H.Z conceived the study and designed the experiments. T.H.Z. and L.D. performed the experiments. L.D., T.H.Z., J.L.S., and R.S analyzed the data. J.P.B. performed the analysis of Potts model. T.H.Z. and L.D. wrote the manuscript with inputs from all authors.

## Acknowledgement

We thank UCLA/CFAR Virology Core Lab for doing p24 ELISA. Work in the Virology Core Lab was supported by UCLA CFAR grant 5P30 AI028697.

---

**Figure S1.**
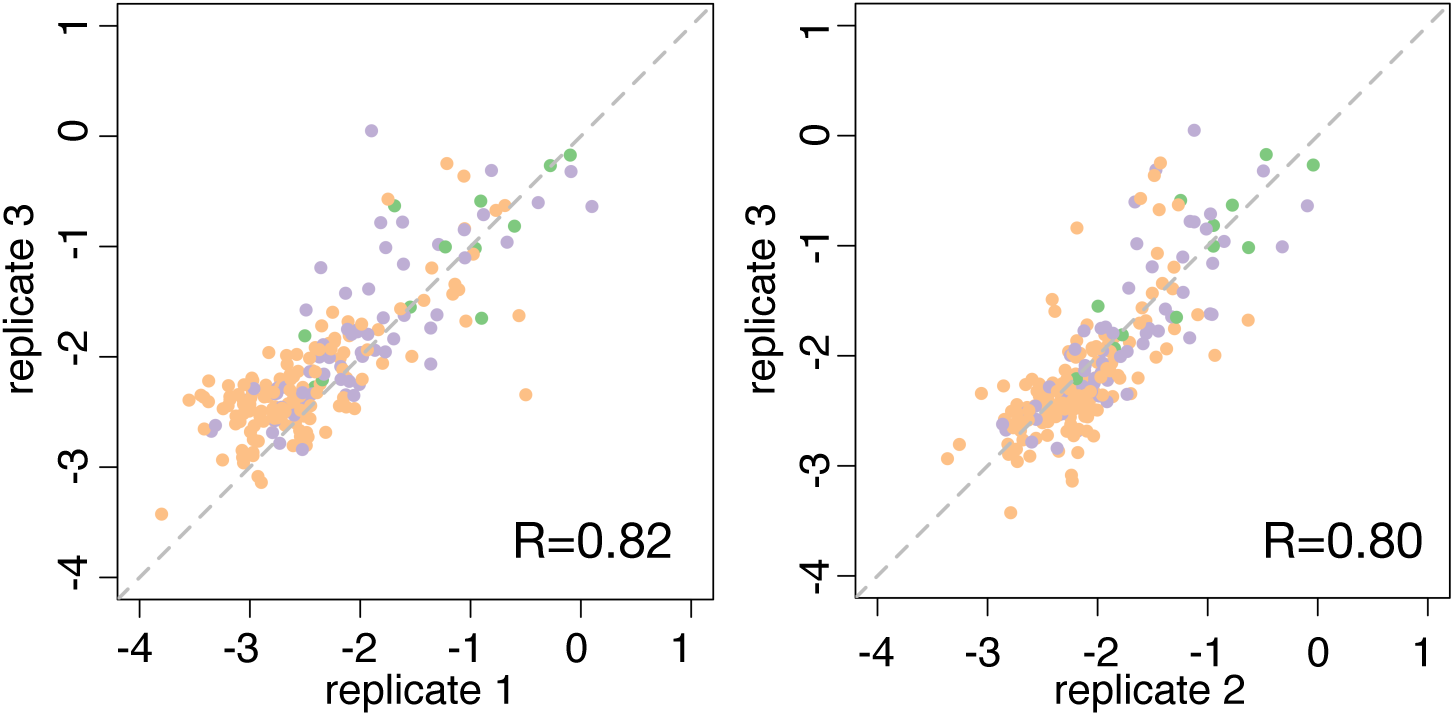
The correlation of relative fitness among biological replicates. All single mutants, double mutants and triple mutants are shown. R stands for Pearson correlation coefficient.

**Figure S2.**
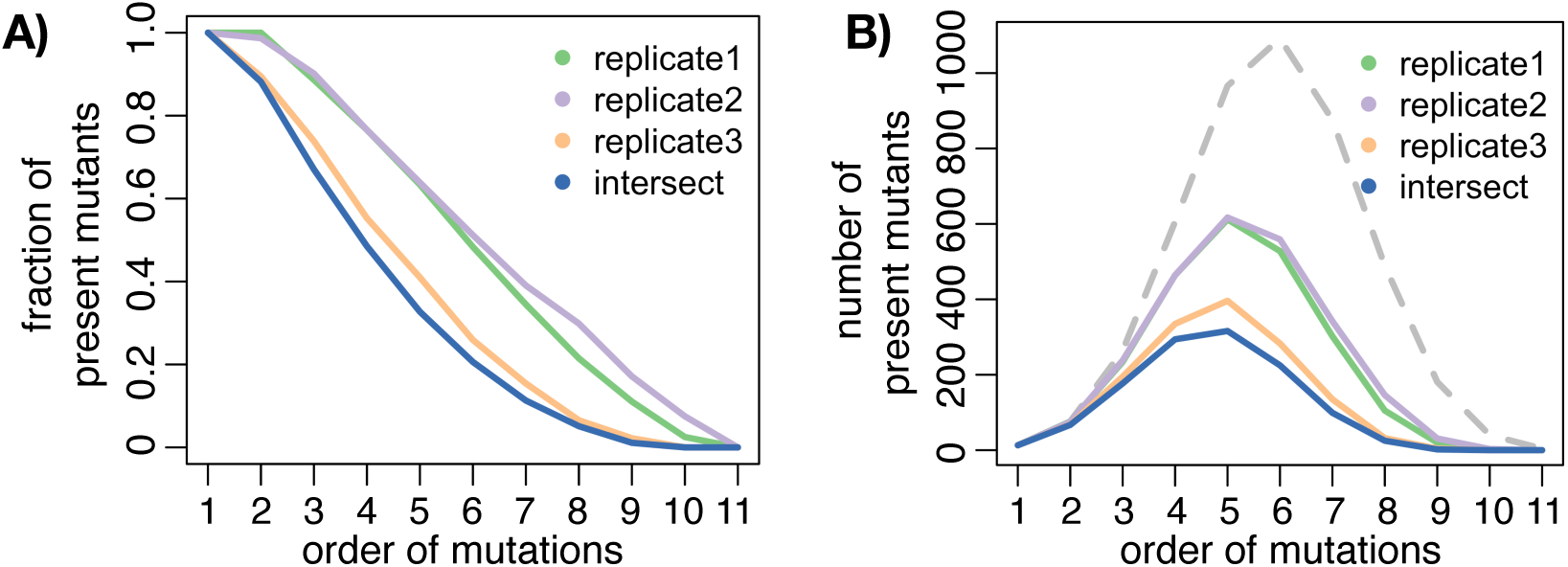
Coverage of protease mutant library. A) Fraction of expected protease mutants in each transfection virus library. B) Number of mutant in each transfection virus library. Dashed line represents the number of all possible combinations of mutations.

**Figure S3.**
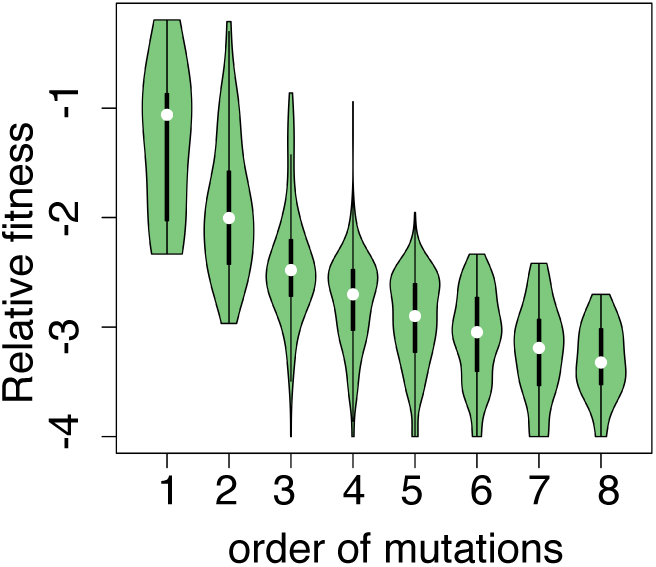
Relative fitness of different order of mutations.

**Figure S4.**
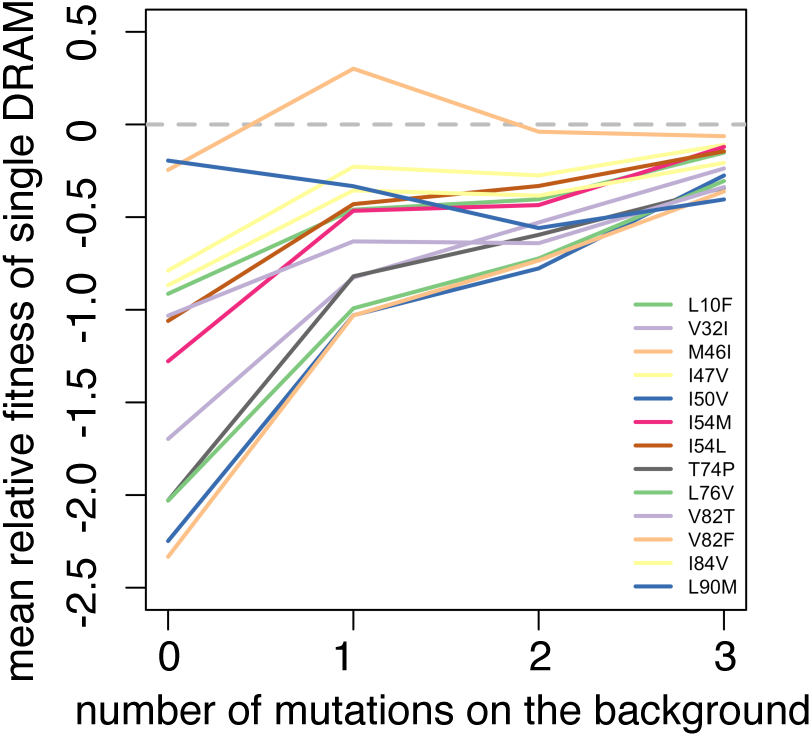
Relative fitness of single DRAMs on different genetic backgrounds.

**Figure S5.**
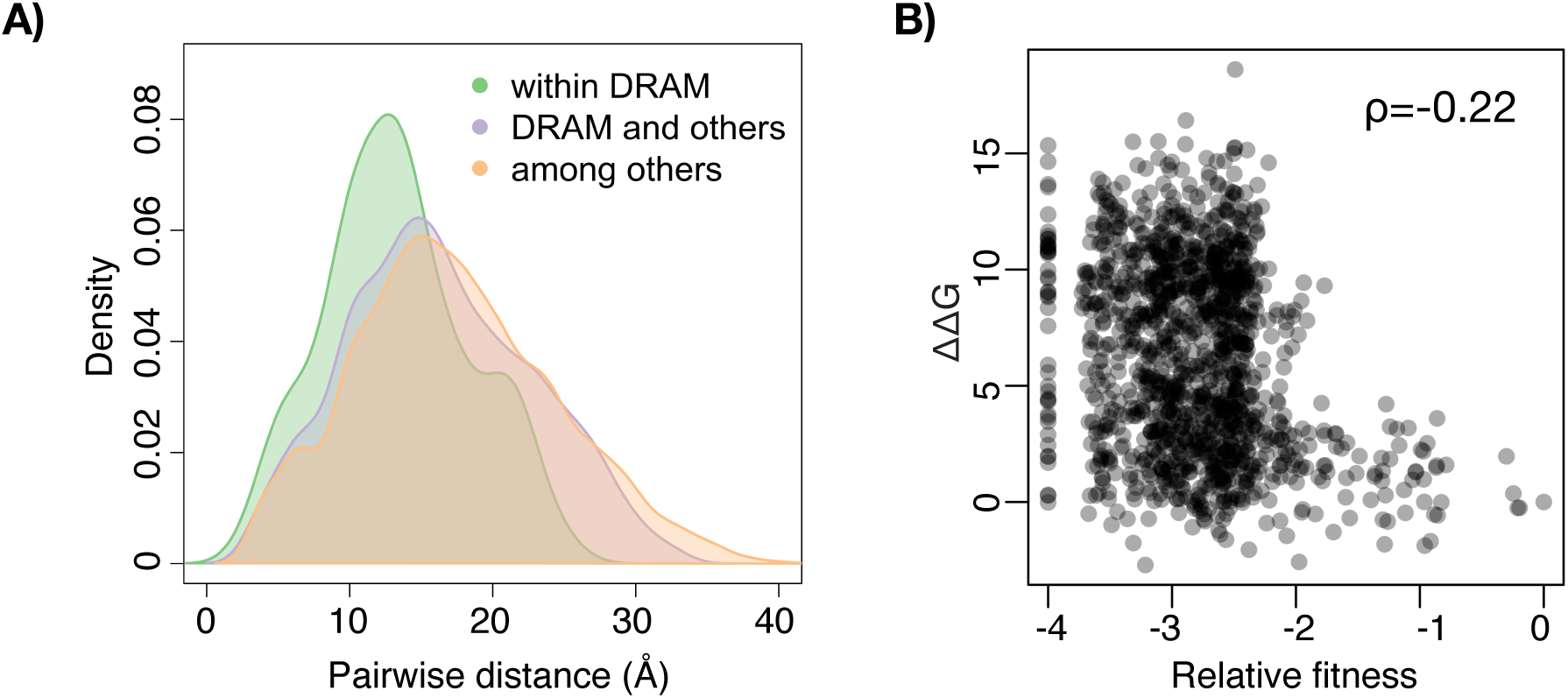
Structure insights on drug resistance associated mutations. A) Distribution of pairwise distance among drug resistance associated residues and other residues. B) Correlation between mutants’ relative fitness and protein stability (ΔΔG). ΔΔG is predicted by FoldX. ρ stands for Spearman’s correlation coefficient.

