## Supplementary file 1 for "Predominance of positive epistasis among drug resistance-associated mutations in HIV-1 protease"

**Supplementary Table1: Relative fitness of all mutants in this research.**

| mutants | relative fitness |
|---|---|
| WT | 0 |
| L90M | -0.1947913 |
| M46I_I54M | -0.210517798 |
| M46I | -0.245238013 |
| M46I_I84V | -0.301304513 |
| I47V | -0.788208628 |
| L10F_I47V | -0.826837086 |
| V32I_M46I | -0.856170224 |
| M46I_I47V_I84V | -0.86114999 |
| M46I_I54L | -0.863881775 |
| I84V | -0.867699991 |
| V32I_I47V | -0.883212633 |
| M46I_I47V_I50V_I54M_L76V_V82T_L90M | -0.883212633 |
| L10F | -0.913817233 |
| V32I_M46I_I47V_I54L | -0.940149914 |
| L10F_V32I_I47V | -0.960941838 |
| V32I_M46I_I47V | -0.963964395 |
| V32I_I47V_I54L | -0.968889717 |
| M46I_I47V | -0.972647054 |
| M46I_T74P | -0.990364837 |
| V82T | -1.032011112 |
| I84V_L90M | -1.033638197 |
| I54L | -1.060154581 |
| M46I_I54L_T74P | -1.092675437 |
| M46I_I54M_L90M | -1.116853449 |
| M46I_L90M | -1.126921682 |
| M46I_I54L_V82T | -1.167354998 |
| I47V_I84V | -1.184826166 |
| V32I_I47V_I50V_L76V_V82T_I84V_L90M | -1.184826166 |
| M46I_V82T | -1.240289455 |
| I47V_I54L | -1.240961403 |
| V32I_M46I_I47V_I54M | -1.259988096 |
| M46I_I47V_I54L | -1.272026076 |
| I54M | -1.277808526 |
| V32I_I47V_I54M | -1.284242454 |
| V32I_I47V_L90M | -1.298435 |
| I47V_L90M | -1.300364322 |
| I54L_L90M | -1.305172205 |
| M46I_I84V_L90M | -1.309589752 |
| M46I_I54L_L90M | -1.360177638 |
| V32I_I47V_I84V | -1.365355229 |

| | |
|---|---|
| I47V_I54M | -1.395415771 |
| M46I_I54M_V82T | -1.487544746 |
| M46I_I54M_I84V | -1.513918186 |
| I54M_L90M | -1.565735772 |
| V82T_I84V | -1.588215883 |
| V82T_L90M | -1.593221873 |
| V32I_M46I_I54L | -1.598126276 |
| V82T_I84V_L90M | -1.630568215 |
| I54L_V82T | -1.675293989 |
| M46I_I50V | -1.675709393 |
| I47V_V82T | -1.684426388 |
| V32I | -1.697353755 |
| V82F_I84V_L90M | -1.748461422 |
| V82F_L90M | -1.761803237 |
| T74P_V82T | -1.768231494 |
| L76V_V82T_I84V | -1.769719912 |
| M46I_I54L_I84V | -1.774813901 |
| V82F_I84V | -1.775726323 |
| M46I_I47V_I54M | -1.784561117 |
| I47V_I50V | -1.794096798 |
| I54M_V82F | -1.816938041 |
| V32I_M46I_I47V_L90M | -1.843399904 |
| I54L_T74P | -1.893931642 |
| T74P_I84V | -1.894488548 |
| M46I_L76V | -1.91093759 |
| V32I_I47V_I54L_L90M | -1.91882379 |
| V32I_I47V_I54M_T74P_V82F | -1.934522891 |
| L76V_V82T | -1.937497084 |
| V32I_I54L | -1.944788899 |
| T74P_V82F_I84V | -1.947423102 |
| V32I_M46I_I54M | -1.953058489 |
| V32I_M46I_I47V_I54L_L90M | -1.954856554 |
| L76V_V82F_I84V | -1.957625437 |
| V32I_I50V_I54M_L76V_V82T | -1.95818681 |
| I47V_I54L_T74P | -1.96049561 |
| V32I_M46I_I47V_I84V | -1.968577704 |
| V32I_M46I_I50V | -1.970550543 |
| V32I_I47V_I54L_T74P_I84V | -1.972110531 |
| L10F_V32I | -1.974487084 |
| L76V_V82T_L90M | -1.980235117 |
| M46I_I47V_I54L_T74P_L76V | -1.983374356 |
| V32I_M46I_I84V | -1.9946296 |
| V32I_I47V_T74P_V82F_I84V | -2.000519279 |

| | |
|---|---|
| I54M_I84V | -2.004383013 |
| I50V_T74P | -2.011847712 |
| L76V_V82F_I84V_L90M | -2.011943547 |
| M46I_I47V_I50V_I54M | -2.01525405 |
| I47V_I50V_I54M | -2.024613231 |
| L76V_V82F | -2.026035294 |
| T74P | -2.029118972 |
| L76V | -2.030416901 |
| M46I_I50V_I54L_L76V_L90M | -2.030737477 |
| I54M_V82T | -2.046764597 |
| I47V_T74P_I84V | -2.052456726 |
| I47V_I84V_L90M | -2.05493939 |
| V32I_I47V_I50V | -2.063303705 |
| T74P_L76V_I84V | -2.065125421 |
| L76V_I84V | -2.065693468 |
| V32I_I47V_T74P_L76V | -2.073184427 |
| V32I_L90M | -2.074073617 |
| V32I_M46I_T74P | -2.075673277 |
| M46I_I50V_I54M_V82F_I84V_L90M | -2.076494967 |
| T74P_L76V | -2.077428745 |
| M46I_I54L_L76V | -2.078195193 |
| M46I_I47V_L90M | -2.081019043 |
| V32I_I47V_I50V_I54L_L76V_V82T | -2.087512327 |
| M46I_I47V_I50V_I54L | -2.090353645 |
| I54L_T74P_I84V | -2.09563053 |
| M46I_V82F | -2.096463289 |
| I47V_I50V_I54L | -2.097764705 |
| M46I_I47V_I84V_L90M | -2.099780257 |
| M46I_I50V_I54M | -2.112316444 |
| I47V_I50V_I54M_L76V_I84V | -2.113316378 |
| V32I_M46I_I47V_I50V | -2.118612048 |
| V32I_M46I_I50V_I54M | -2.121769552 |
| M46I_I54M_T74P | -2.124972748 |
| I50V_I54M | -2.125932254 |
| M46I_V82T_I84V | -2.128404194 |
| V32I_M46I_L90M | -2.128509809 |
| V32I_M46I_I47V_I50V_I54L | -2.129531748 |
| M46I_I54M_V82F | -2.130902562 |
| I54L_I84V | -2.135465128 |
| M46I_I47V_I54L_T74P | -2.140006582 |
| M46I_I47V_I50V | -2.144028864 |
| I47V_I54L_L90M | -2.148680321 |
| I50V_I54L | -2.151917199 |

| | |
|---|---|
| L76V_V82T_I84V_L90M | -2.152845028 |
| V32I_I50V_I54L | -2.153086767 |
| V32I_I47V_I54M_T74P_L76V | -2.154680046 |
| L76V_V82F_L90M | -2.154717874 |
| I47V_I50V_I54M_L76V_V82F | -2.160834114 |
| T74P_L76V_V82F_I84V | -2.175853194 |
| V32I_M46I_I47V_I50V_I54M | -2.184572229 |
| V32I_I54L_L76V_V82T_L90M | -2.189795396 |
| I54L_V82F | -2.190232595 |
| I54L_T74P_L76V_V82F | -2.195222292 |
| M46I_I47V_I50V_L76V_I84V_L90M | -2.195222292 |
| M46I_I50V_I54L | -2.198939091 |
| V32I_I50V_I54L_L76V_V82F_I84V_L90M | -2.199340714 |
| I54M_V82T_I84V | -2.200659632 |
| M46I_I50V_I54L_L76V_V82F_I84V_L90M | -2.205589664 |
| M46I_I47V_I50V_I54M_V82T_I84V | -2.207218347 |
| V32I_I47V_T74P_V82T | -2.207973642 |
| I47V_I50V_I54M_L76V_V82T_L90M | -2.208680741 |
| V32I_T74P_L76V_V82F | -2.211487126 |
| V32I_M46I_L76V_V82F_L90M | -2.21426045 |
| V32I_I47V_I50V_I54M | -2.215570595 |
| M46I_I54L_L76V_I84V | -2.218914861 |
| M46I_I50V_I54L_L76V | -2.220901896 |
| L76V_I84V_L90M | -2.225027794 |
| V32I_M46I_I50V_I54L | -2.234284901 |
| V32I_I54M | -2.236784796 |
| M46I_I54L_V82F | -2.237345333 |
| M46I_I47V_I54M_V82F | -2.23933252 |
| M46I_I47V_I50V_I54M_I84V | -2.243136907 |
| M46I_I47V_I54L_L90M | -2.247681386 |
| I50V | -2.248032685 |
| V32I_I50V_I54M_L76V_V82T_I84V | -2.249494475 |
| V32I_I54M_T74P | -2.250094164 |
| V32I_I47V_I50V_I54L | -2.250888464 |
| I54L_T74P_V82F | -2.250930956 |
| I47V_L76V_V82T_I84V_L90M | -2.252586226 |
| I47V_I54M_V82T_I84V | -2.257536251 |
| I54L_L76V_V82F | -2.260651203 |
| M46I_I47V_I50V_I54M_L76V_V82T_I84V | -2.261510143 |
| V32I_M46I_I50V_I54M_V82T_I84V_L90M | -2.261525606 |
| V32I_M46I_I47V_I54M_L76V_V82T_I84V_L90M | -2.263581611 |
| I47V_I50V_I54M_L76V_V82F_I84V_L90M | -2.265263792 |
| M46I_I47V_I50V_V82T | -2.266425396 |

| | |
|---|---|
| V32I_M46I_I47V_I50V_I54M_V82T_I84V | -2.268739886 |
| M46I_I47V_I54L_L76V_V82T_L90M | -2.268976643 |
| V32I_M46I_L76V_I84V_L90M | -2.268976643 |
| V32I_M46I_I47V_I50V_I54M_L76V_I84V_L90M | -2.270464986 |
| I54M_L76V | -2.271981867 |
| M46I_I47V_I54M_L76V_V82F_I84V | -2.274143518 |
| I47V_I50V_I54L_L76V | -2.275724356 |
| V32I_I50V_I54M_V82T | -2.275898962 |
| V32I_M46I_I50V_I54M_L76V_V82F | -2.276697154 |
| V32I_I47V_I50V_I54M_L76V_I84V_L90M | -2.27730441 |
| V32I_I50V_T74P_V82F | -2.27730441 |
| V32I_I47V_V82F | -2.277491332 |
| M46I_I54M_T74P_V82F_I84V | -2.279334536 |
| V32I_M46I_I47V_I50V_I54L_V82F | -2.280390425 |
| V32I_I47V_I50V_I54M_V82F_I84V | -2.280471435 |
| V32I_M46I_I47V_I54M_L76V_V82T_I84V | -2.282337614 |
| I47V_I50V_I54L_I84V_L90M | -2.283543603 |
| V32I_M46I_I54M_L76V_V82F_L90M | -2.28477091 |
| V32I_M46I_I47V_I50V_I54L_L76V_V82F_L90M | -2.28477091 |
| I54M_V82F_I84V | -2.286755528 |
| M46I_I47V_I54L_T74P_V82F_I84V | -2.289125129 |
| M46I_I47V_I50V_L76V_V82F | -2.290375965 |
| I50V_I54L_L76V_V82T_I84V | -2.291086332 |
| I47V_I50V_I54M_L76V_V82T | -2.292557286 |
| M46I_V82T_L90M | -2.293155454 |
| V32I_M46I_I47V_I50V_I54L_V82T_L90M | -2.298493709 |
| I50V_I54M_V82T_I84V | -2.299870024 |
| V32I_I50V | -2.301135711 |
| I54L_T74P_L76V_V82F_I84V | -2.301381204 |
| V32I_I47V_I50V_I54M_I84V | -2.302771743 |
| M46I_I47V_I50V_V82T_L90M | -2.304974296 |
| I50V_I54M_L76V_V82F | -2.306160533 |
| M46I_I47V_I54M_L76V_V82T_L90M | -2.308904589 |
| I50V_I54L_L76V_I84V | -2.310403463 |
| M46I_I50V_L76V_V82T | -2.31128038 |
| V32I_I50V_I54L_V82F_L90M | -2.314734133 |
| M46I_I50V_I54L_L76V_V82T_L90M | -2.314734133 |
| I47V_I54L_L76V | -2.315991912 |
| M46I_I47V_I50V_I54M_V82F_L90M | -2.316200925 |
| I50V_L76V_V82F | -2.317473019 |
| I50V_I54M_V82F_I84V | -2.317647413 |
| I47V_I54M_L76V_V82T | -2.319141151 |
| M46I_I47V_V82T | -2.319272509 |

| | |
|---|---|
| I50V_L76V_V82T_I84V | -2.320159252 |
| M46I_I47V_I50V_I54L_V82F_I84V | -2.32362034 |
| L10F_V32I_I47V_T74P | -2.324279451 |
| I47V_I50V_I54M_L76V | -2.325858128 |
| V32I_I50V_I54L_L76V_V82T | -2.328715752 |
| M46I_T74P_L76V | -2.32980854 |
| M46I_I47V_I50V_I54M_V82F_I84V_L90M | -2.330649852 |
| I47V_I50V_I54M_L76V_I84V_L90M | -2.332190182 |
| M46I_I47V_I50V_I54L_L76V_V82T_I84V_L90M | -2.3326093 |
| V82F | -2.333144788 |
| M46I_I47V_I50V_I54L_L76V_V82F | -2.333911756 |
| V32I_I54L_T74P | -2.334207993 |
| I47V_I50V_I54M_L76V_V82T_I84V | -2.336818692 |
| M46I_I50V_I54M_V82F_I84V | -2.337577767 |
| M46I_I50V_V82F_I84V | -2.337957179 |
| V32I_I54M_L76V_V82F_I84V | -2.338846389 |
| I50V_I54M_L76V | -2.339112335 |
| I54M_L76V_V82F_I84V | -2.341075762 |
| V32I_I47V_T74P_I84V | -2.341811338 |
| V32I_I50V_I54M_V82T_I84V | -2.341876499 |
| V32I_M46I_I47V_I50V_V82T_I84V | -2.341987361 |
| I54L_V82F_I84V | -2.344009083 |
| M46I_I47V_I50V_L76V | -2.34639488 |
| V32I_I84V | -2.347071087 |
| V32I_M46I_I47V_T74P | -2.347452121 |
| M46I_I47V_I50V_I54M_V82F | -2.348741097 |
| M46I_I50V_L76V_V82F_I84V | -2.349634354 |
| I47V_T74P_V82F_L90M | -2.349634354 |
| V32I_M46I_I50V_I84V | -2.351223238 |
| V32I_I50V_I54L_L76V_V82T_I84V | -2.352548481 |
| M46I_I50V_I54M_L76V_I84V | -2.355129003 |
| I47V_I54M_V82T | -2.35520235 |
| V32I_M46I_I47V_I54L_I84V | -2.356771227 |
| V32I_I50V_I54L_I84V | -2.357479022 |
| V32I_M46I_I50V_I54M_V82F | -2.357712128 |
| V32I_M46I_I47V_V82T | -2.359169697 |
| I47V_T74P_V82T | -2.359179352 |
| L10F_I47V_T74P_V82F | -2.361359345 |
| V32I_I47V_I50V_I54M_L90M | -2.361855331 |
| I47V_I50V_L76V_V82T | -2.361919566 |
| V32I_I50V_L76V_V82F_L90M | -2.363219039 |
| V32I_M46I_I50V_I54L_L76V_V82F_L90M | -2.363219039 |
| M46I_I50V_I54M_V82T_I84V | -2.363291175 |

| | |
|---|---|
| I54L_L76V | -2.364017665 |
| I47V_I50V_I54L_T74P_V82F | -2.364845434 |
| I50V_I54M_V82T | -2.365122999 |
| V32I_I47V_I50V_I54M_L76V_V82T_I84V | -2.36547885 |
| V32I_M46I_I54L_L76V_V82T | -2.365760643 |
| V32I_M46I_I47V_I54M_L76V_I84V | -2.367345268 |
| I54L_V82F_L90M | -2.368218798 |
| V32I_L76V_V82T_I84V_L90M | -2.369091795 |
| I54M_L76V_V82T | -2.369862785 |
| V32I_I47V_I50V_I54L_L76V_L90M | -2.371491565 |
| V32I_I47V_I50V_I54M_L76V_V82F | -2.373371675 |
| V32I_M46I_I50V_L76V_I84V | -2.374609148 |
| I50V_I54M_V82F_L90M | -2.375431973 |
| M46I_I47V_V82T_I84V | -2.378567209 |
| V32I_I47V_I54M_L90M | -2.379582294 |
| I47V_I54L_T74P_L76V | -2.380097931 |
| M46I_I54M_L76V_V82F | -2.381294113 |
| M46I_I50V_V82T | -2.381780175 |
| M46I_I50V_I54M_L76V_V82T_I84V | -2.382217344 |
| I50V_I54L_I84V_L90M | -2.382247759 |
| I47V_I50V_I54M_V82F_L90M | -2.383028397 |
| I47V_I50V_I54M_V82T | -2.383360443 |
| M46I_I47V_V82F | -2.383886663 |
| V32I_I54M_L76V_V82T_I84V | -2.385899689 |
| M46I_V82F_L90M | -2.387089079 |
| V32I_M46I_I47V_I54M_V82T | -2.388727512 |
| M46I_I47V_I50V_I54M_V82T_I84V_L90M | -2.390307757 |
| M46I_I47V_I54M_V82T | -2.390537357 |
| M46I_I47V_I54M_L90M | -2.390655471 |
| V32I_I47V_I54M_L76V | -2.391283596 |
| I47V_I54L_L76V_V82F | -2.392832715 |
| V32I_M46I_I47V_I50V_I54L_L76V_L90M | -2.393915379 |
| M46I_I47V_I50V_I54L_L76V_V82T | -2.394280488 |
| V32I_M46I_I50V_V82T_L90M | -2.394280488 |
| M46I_I54M_I84V_L90M | -2.394639216 |
| M46I_I54M_L76V | -2.395792727 |
| V32I_M46I_I47V_I50V_I54M_V82F_I84V | -2.396383573 |
| M46I_I47V_I54L_I84V | -2.397339377 |
| V32I_I47V_V82T | -2.398296166 |
| V32I_I47V_I50V_I54M_V82T_I84V_L90M | -2.398296166 |
| M46I_I50V_I54M_L76V | -2.398645254 |
| V32I_M46I_I47V_L76V_V82T | -2.398853644 |
| I50V_I54L_V82T_I84V | -2.401538141 |

| | |
|---|---|
| M46I_I50V_I54L_V82F_I84V | -2.401661833 |
| V32I_M46I_V82T | -2.4021392 |
| V32I_I47V_I54M_V82T_I84V | -2.402518654 |
| M46I_I50V_I54L_V82T_I84V | -2.402988154 |
| I54M_V82T_I84V_L90M | -2.403275624 |
| I54L_V82T_I84V | -2.404548498 |
| V32I_I50V_I54M | -2.405229862 |
| V32I_I47V_T74P_V82F | -2.405311639 |
| V32I_M46I_I47V_I50V_I54L_V82T | -2.405498295 |
| M46I_I47V_I54M_L76V | -2.405757239 |
| V32I_M46I_I54L_V82T | -2.405815487 |
| I47V_V82F | -2.406213832 |
| V32I_I47V_I50V_I54L_T74P | -2.407493516 |
| V32I_M46I_I50V_I54L_L76V_V82T | -2.407697082 |
| M46I_I54L_V82T_L90M | -2.407814859 |
| V32I_I50V_L76V_I84V | -2.409103819 |
| V32I_M46I_L76V_V82T_I84V | -2.409641162 |
| V32I_M46I_I47V_I50V_L76V_V82T_I84V_L90M | -2.410643689 |
| I50V_I54L_L90M | -2.414098887 |
| V32I_I47V_I50V_L76V_I84V_L90M | -2.415721154 |
| M46I_I47V_I54L_L76V | -2.416415171 |
| V32I_M46I_I50V_L76V_V82F | -2.416639034 |
| M46I_I50V_I54M_I84V_L90M | -2.417396475 |
| V32I_I50V_V82T_I84V_L90M | -2.417396475 |
| V32I_M46I_I47V_I50V_I54M_L76V_I84V | -2.417667199 |
| V32I_M46I_I47V_I50V_I54L_L76V | -2.418042487 |
| V32I_M46I_I50V_I54M_L76V_V82F_I84V | -2.421667387 |
| I47V_I54L_T74P_V82F_I84V | -2.421842846 |
| M46I_L76V_V82F | -2.423097791 |
| I54L_L76V_V82T_I84V | -2.424136079 |
| V32I_M46I_I50V_I54M_V82T_I84V | -2.424164003 |
| V32I_I47V_I54L_L76V | -2.424359354 |
| V32I_M46I_I47V_I50V_I54L_V82T_I84V | -2.42443583 |
| M46I_I47V_T74P_L76V | -2.425980383 |
| V32I_M46I_I50V_V82T_I84V | -2.426176418 |
| V32I_M46I_I47V_I50V_I54M_L90M | -2.428630303 |
| V32I_M46I_I47V_L76V_I84V_L90M | -2.428677485 |
| M46I_I47V_I50V_T74P | -2.430117374 |
| V32I_M46I_I50V_L76V_V82T_I84V | -2.430611508 |
| M46I_I47V_I50V_I54M_L76V | -2.431830587 |
| V32I_M46I_I54M_V82T | -2.432561039 |
| V32I_M46I_I47V_I50V_L76V_V82T_I84V | -2.432633882 |
| M46I_I47V_I50V_I54M_V82T | -2.432791394 |

| | |
|---|---|
| I47V_I50V_I54M_I84V | -2.433174885 |
| V32I_I54M_L76V_V82T | -2.433265549 |
| I54L_L76V_V82F_I84V | -2.433486688 |
| I47V_I54M_V82F | -2.433995889 |
| V32I_I47V_I50V_L76V_V82T_L90M | -2.433995889 |
| V32I_M46I_I50V_I54L_V82T_L90M | -2.434250174 |
| V32I_I54M_V82T | -2.434408493 |
| I47V_I54M_T74P_V82F_I84V | -2.435752929 |
| M46I_L76V_V82T_I84V_L90M | -2.4361092 |
| V32I_M46I_I50V_I54L_L76V | -2.436156659 |
| V32I_I47V_I54M_V82F_L90M | -2.436652377 |
| V32I_M46I_I50V_I54L_L76V_V82F | -2.438209814 |
| I47V_I50V_V82F_I84V | -2.438290002 |
| M46I_V82F_I84V | -2.439448051 |
| V32I_I47V_I50V_I54L_L76V | -2.439582498 |
| M46I_I47V_L76V_I84V_L90M | -2.440077483 |
| I47V_I54L_L76V_V82T_I84V | -2.440338393 |
| M46I_L76V_I84V | -2.44102732 |
| V32I_M46I_I47V_V82T_I84V_L90M | -2.441127064 |
| V32I_M46I_I47V_I50V_I54M_V82T_L90M | -2.441870157 |
| V32I_I47V_I54L_V82T | -2.442752089 |
| M46I_I47V_L76V | -2.443753615 |
| I54M_L76V_V82F | -2.443897669 |
| I47V_I54L_L76V_I84V | -2.443946569 |
| V32I_M46I_I50V_L76V | -2.445344639 |
| I47V_I50V_I54M_V82T_I84V_L90M | -2.445344639 |
| L76V_L90M | -2.447110573 |
| I47V_I54L_V82F_I84V | -2.447591062 |
| M46I_I50V_I54M_I84V | -2.448362347 |
| I54M_T74P | -2.449231785 |
| M46I_I50V_I54M_V82T_I84V_L90M | -2.449888743 |
| V32I_I50V_L76V_V82F_I84V_L90M | -2.449900961 |
| M46I_I47V_I54L_L76V_V82T | -2.450362888 |
| V32I_M46I_I54L_V82F | -2.451773014 |
| V32I_M46I_L76V_I84V | -2.451926312 |
| V32I_I47V_I50V_L76V | -2.453911178 |
| M46I_I47V_I50V_V82T_I84V | -2.454010592 |
| M46I_I50V_I54M_V82F | -2.454295754 |
| I47V_I50V_I54M_V82F | -2.45434637 |
| M46I_I50V_L76V_V82T_I84V | -2.454376152 |
| V32I_I47V_I54L_I84V | -2.456190988 |
| I47V_I54M_L76V | -2.456577033 |
| V32I_M46I_I50V_I54M_I84V | -2.45702118 |

| | |
|---|---|
| I47V_I54M_L76V_V82F_I84V | -2.457600735 |
| I47V_I50V_V82T | -2.458067729 |
| I47V_I54M_L76V_V82F | -2.459263944 |
| M46I_I54L_V82T_I84V_L90M | -2.460129052 |
| I47V_I54M_I84V | -2.462173637 |
| I47V_I50V_I84V | -2.462821124 |
| I47V_T74P_L76V | -2.463201117 |
| V32I_I47V_I50V_I54L_L76V_V82F | -2.464186587 |
| I47V_I54M_L90M | -2.464289912 |
| V32I_M46I_I50V_V82T_I84V_L90M | -2.464289912 |
| V32I_M46I_I47V_I54L_L76V | -2.467518058 |
| I50V_L76V | -2.468129921 |
| V32I_M46I_I54M_L76V_V82T_I84V | -2.468243328 |
| I47V_L76V_V82T | -2.468284684 |
| I47V_I50V_L76V | -2.468322756 |
| V32I_T74P | -2.468495855 |
| M46I_I50V_V82F_I84V_L90M | -2.469134616 |
| I50V_I54L_L76V | -2.469575914 |
| I50V_I54L_V82F_I84V | -2.470235279 |
| V32I_I50V_I54M_V82F_L90M | -2.470235279 |
| M46I_I50V_I54L_V82T_I84V_L90M | -2.470293388 |
| M46I_I54L_V82F_I84V | -2.470435615 |
| V32I_L76V_V82F_L90M | -2.471385497 |
| I54L_T74P_L76V | -2.472321481 |
| M46I_I54M_L76V_V82T | -2.472938816 |
| I47V_I50V_I54M_V82T_I84V | -2.474363332 |
| I50V_I54L_V82F | -2.47453636 |
| M46I_I47V_I54L_L76V_V82F_I84V | -2.475027274 |
| M46I_I47V_I50V_I54M_L76V_V82F_I84V_L90M | -2.47688319 |
| I50V_I54M_I84V_L90M | -2.476956936 |
| V32I_M46I_I47V_I54L_V82T_I84V | -2.477140119 |
| V32I_M46I_I54M_V82T_I84V | -2.4775001 |
| V32I_L76V_I84V | -2.478646235 |
| I47V_T74P_V82F | -2.479433262 |
| V32I_I47V_I54M_L76V_V82F | -2.479733117 |
| M46I_I54L_L76V_I84V_L90M | -2.480395081 |
| I50V_I54L_L76V_I84V_L90M | -2.481065555 |
| M46I_I54M_L76V_V82T_I84V | -2.481343227 |
| M46I_I47V_L76V_I84V | -2.482412078 |
| M46I_I50V_L76V_V82F_L90M | -2.483135601 |
| V32I_M46I_L76V_V82F_I84V | -2.483239809 |
| V32I_M46I_I47V_I50V_I54M_L76V_V82T_I84V | -2.48356517 |
| M46I_I47V_I50V_I54L_L76V | -2.483574992 |

| | |
|---|---|
| I47V_V82F_I84V | -2.48367921 |
| I47V_L76V | -2.484188188 |
| I54M_L76V_V82T_I84V | -2.484987464 |
| V32I_I47V_I50V_L76V_V82T | -2.485003877 |
| V32I_M46I_I47V_I50V_I54L_L90M | -2.485275 |
| V32I_M46I_I47V_I54M_L76V_V82F_L90M | -2.485275 |
| V32I_I47V_V82T_I84V | -2.486077956 |
| I50V_I54M_L76V_I84V | -2.486266218 |
| V32I_I47V_I50V_I54L_V82F | -2.48727376 |
| V32I_M46I_I47V_I50V_L76V | -2.488230508 |
| I47V_T74P | -2.489969406 |
| M46I_I47V_I50V_I54L_L76V_V82T_I84V | -2.490374158 |
| V32I_M46I_I47V_I50V_I54L_L76V_V82T | -2.490879111 |
| I47V_I54L_V82F | -2.49101853 |
| M46I_I50V_I54M_V82T | -2.491070322 |
| M46I_I47V_I50V_V82T_I84V_L90M | -2.491569564 |
| V32I_I47V_I50V_I54L_L76V_V82T_L90M | -2.491569564 |
| M46I_I47V_I50V_I54M_L76V_V82F_L90M | -2.491569564 |
| M46I_I54L_L76V_V82T_L90M | -2.491615516 |
| I47V_I50V_I54L_L76V_V82T_I84V | -2.491712436 |
| V32I_I47V_I54M_L76V_I84V | -2.491903495 |
| V32I_I47V_I50V_I54L_V82F_L90M | -2.491903495 |
| M46I_I47V_I54M_L76V_L90M | -2.492051578 |
| V32I_M46I_L76V | -2.493082809 |
| V32I_T74P_V82T | -2.493082809 |
| M46I_T74P_V82F_I84V | -2.493217748 |
| M46I_I50V_I54L_L90M | -2.494376896 |
| V32I_M46I_I50V_I54L_V82T | -2.494637077 |
| V32I_I47V_I50V_V82T_I84V | -2.495639742 |
| M46I_I47V_I50V_I54L_V82T | -2.4964324 |
| V32I_M46I_I47V_I50V_V82F | -2.496780727 |
| V32I_I50V_L76V_V82F_I84V | -2.496816974 |
| M46I_I47V_I50V_I54L_L76V_I84V | -2.49714855 |
| I47V_I50V_I54L_L76V_V82T | -2.497468813 |
| M46I_I47V_I54L_V82T | -2.497898144 |
| M46I_I50V_I54M_L76V_V82F_I84V_L90M | -2.497898144 |
| I54L_L76V_L90M | -2.498534066 |
| I47V_L76V_V82F_I84V | -2.499142988 |
| V32I_I47V_V82T_I84V_L90M | -2.499360063 |
| V32I_M46I_I50V_I54M_L76V_V82T_I84V | -2.49938389 |
| I47V_I50V_I54M_V82F_I84V | -2.499774127 |
| V32I_I50V_L76V_V82T_I84V | -2.500351209 |
| V32I_M46I_I47V_I50V_I54M_L76V_V82T | -2.50037071 |

| | |
|---|---|
| V32I_T74P_V82F_I84V | -2.50037071 |
| M46I_I47V_I50V_V82F | -2.501301221 |
| M46I_I54M_V82T_I84V_L90M | -2.501565948 |
| M46I_I47V_I50V_L76V_V82F_I84V | -2.501908649 |
| V32I_M46I_I50V_L76V_V82F_I84V | -2.502526891 |
| M46I_I47V_I50V_I54L_L76V_V82F_I84V_L90M | -2.503070519 |
| V32I_M46I_L76V_V82T_I84V_L90M | -2.503085497 |
| M46I_I47V_I50V_I54L_L76V_V82F_I84V | -2.503900422 |
| V32I_M46I_I47V_I50V_I84V_L90M | -2.503900422 |
| V32I_I50V_I54M_L76V_I84V | -2.504090893 |
| V32I_M46I_V82T_I84V_L90M | -2.504548192 |
| V32I_I47V_L76V | -2.504571884 |
| M46I_I47V_I54M_L76V_I84V | -2.505166068 |
| V32I_M46I_I50V_V82F_L90M | -2.505166068 |
| V32I_M46I_I47V_I50V_L76V_I84V_L90M | -2.505166068 |
| I50V_I54L_V82T | -2.505981621 |
| V32I_I54L_V82F_I84V | -2.506277322 |
| V32I_M46I_I54L_L76V | -2.50633531 |
| M46I_I47V_I50V_I54M_L90M | -2.508409062 |
| V32I_M46I_I50V_I54L_L76V_V82F_I84V | -2.509371631 |
| V32I_M46I_I47V_I50V_I54M_L76V_L90M | -2.509710736 |
| V32I_I47V_L76V_I84V | -2.509770305 |
| M46I_L76V_V82F_I84V | -2.510016784 |
| V32I_M46I_L76V_V82F | -2.51013954 |
| M46I_I47V_I54L_L76V_V82T_I84V | -2.51023767 |
| V32I_I47V_I50V_I54M_V82F_L90M | -2.510693508 |
| M46I_V82F_I84V_L90M | -2.511164325 |
| V32I_I54L_T74P_L76V | -2.511164325 |
| M46I_I47V_I54L_V82F | -2.511958694 |
| V32I_M46I_I50V_I54M_L76V_L90M | -2.512453831 |
| M46I_I47V_I54M_V82T_I84V_L90M | -2.512779968 |
| M46I_I47V_I50V_I54L_V82F | -2.513037267 |
| V32I_M46I_I47V_I50V_I54M_L76V_V82F | -2.513079076 |
| V32I_M46I_I50V_V82T | -2.513347747 |
| M46I_I47V_I50V_L76V_L90M | -2.51358096 |
| V32I_I47V_I50V_V82T_I84V_L90M | -2.51446091 |
| V32I_M46I_I54L_V82F_I84V | -2.515018753 |
| M46I_I50V_I54M_L76V_V82F_L90M | -2.515067845 |
| V32I_M46I_I47V_I54M_I84V | -2.515293378 |
| V32I_M46I_I54L_L76V_V82F_I84V | -2.515929494 |
| I50V_I84V | -2.516933342 |
| V32I_I47V_V82F_I84V | -2.517646403 |
| V32I_I50V_I54M_L76V | -2.51821977 |

| | |
|---|---|
| V32I_M46I_I50V_I54M_L90M | -2.519361058 |
| M46I_I47V_V82T_L90M | -2.519941494 |
| V32I_I54M_L76V_V82T_I84V_L90M | -2.520342459 |
| I47V_I50V_I54L_V82T | -2.520981104 |
| I50V_I54L_L76V_V82F | -2.521046894 |
| V32I_I54L_V82T_I84V | -2.521976939 |
| V32I_M46I_V82F_I84V_L90M | -2.522111076 |
| V32I_T74P_I84V | -2.522276959 |
| V32I_M46I_I47V_I50V_I54L_L76V_V82T_L90M | -2.522726224 |
| M46I_I47V_I54M_V82F_I84V | -2.524350421 |
| M46I_I47V_I54L_V82T_I84V | -2.524571493 |
| I50V_I54M_I84V | -2.524601451 |
| I54M_L76V_V82T_I84V_L90M | -2.524751218 |
| I47V_I54M_T74P_V82F | -2.524865623 |
| I50V_I54M_L76V_V82T_I84V | -2.525055559 |
| V32I_M46I_I50V_L76V_L90M | -2.526515149 |
| M46I_I50V_V82F | -2.527083587 |
| V32I_M46I_I47V_V82T_I84V | -2.528482036 |
| V32I_M46I_I50V_I54L_V82F_I84V_L90M | -2.529545702 |
| M46I_I50V_V82T_I84V | -2.529815617 |
| M46I_I47V_I50V_I54M_V82T_L90M | -2.530565469 |
| V32I_I47V_I54M_T74P | -2.531550318 |
| T74P_V82F | -2.53175533 |
| I47V_V82T_I84V | -2.532147614 |
| V32I_I47V_I54M_V82F | -2.533894834 |
| V32I_M46I_V82F_I84V | -2.535958288 |
| V32I_I47V_I50V_I54L_V82F_I84V | -2.536375436 |
| I47V_I50V_I54L_L76V_I84V_L90M | -2.536375436 |
| V32I_I47V_I50V_I54M_T74P_V82F_I84V | -2.536582883 |
| M46I_I47V_I50V_I54M_L76V_V82T_I84V_L90M | -2.536582883 |
| V32I_M46I_I54M_T74P | -2.536582883 |
| V32I_M46I_I47V_I50V_V82T_I84V_L90M | -2.537467351 |
| V32I_I50V_I54M_L76V_V82F_L90M | -2.537872339 |
| V32I_M46I_I47V_V82F | -2.538541287 |
| V32I_I50V_I54L_L76V_V82T_L90M | -2.53905732 |
| V32I_I47V_I50V_I54L_L76V_I84V | -2.540107277 |
| V32I_I47V_I50V_I54M_V82T | -2.540917195 |
| V32I_M46I_I47V_I54L_L76V_V82F | -2.54132064 |
| V32I_I47V_I50V_I54M_V82T_L90M | -2.541700118 |
| V32I_I47V_L76V_V82T_I84V | -2.542396775 |
| M46I_I50V_I84V | -2.54270774 |
| I47V_I50V_V82T_I84V | -2.542907453 |
| I54M_L76V_V82T_L90M | -2.543227027 |

| | |
|---|---|
| V32I_I50V_I54L_L76V_V82F_L90M | -2.543416963 |
| I47V_I54L_I84V | -2.544023814 |
| I54L_L76V_V82T | -2.544095702 |
| I47V_I50V_V82F | -2.545664311 |
| V32I_I54M_L76V_I84V | -2.545788532 |
| M46I_I47V_I54M_L76V_V82T | -2.546612834 |
| V32I_I47V_I54L_V82F_I84V | -2.546675985 |
| V32I_I47V_T74P | -2.547262752 |
| V32I_I47V_I54L_L76V_I84V_L90M | -2.547744368 |
| M46I_I47V_I50V_L76V_I84V | -2.547781945 |
| V32I_M46I_I50V_L76V_V82F_I84V_L90M | -2.548488206 |
| I54L_V82T_I84V_L90M | -2.549159013 |
| V32I_M46I_I54L_I84V | -2.550242846 |
| I50V_I54L_L76V_V82T | -2.550644458 |
| M46I_I47V_L76V_V82F | -2.550863774 |
| M46I_I47V_I50V_I54M_L76V_V82F_I84V | -2.550902769 |
| M46I_I54L_T74P_V82F_I84V | -2.550914457 |
| M46I_I47V_I54M_L76V_V82F_I84V_L90M | -2.551333661 |
| V32I_I47V_I50V_L76V_V82T_I84V | -2.551729174 |
| M46I_I50V_I54L_L76V_I84V | -2.552074811 |
| M46I_L76V_V82F_I84V_L90M | -2.552239168 |
| V32I_M46I_I47V_I50V_I54M_L76V | -2.552378487 |
| V32I_M46I_I47V_I54L_V82F | -2.553154027 |
| V32I_I50V_I54L_V82T_I84V_L90M | -2.553616222 |
| V32I_M46I_I47V_I50V_L76V_I84V | -2.554054132 |
| I50V_L76V_V82T_I84V_L90M | -2.555700714 |
| V32I_M46I_I47V_I50V_L76V_V82T_L90M | -2.555700714 |
| I50V_V82F | -2.555921098 |
| V32I_M46I_I54L_L76V_V82T_I84V | -2.558152802 |
| I54L_T74P_V82F_I84V | -2.558339018 |
| M46I_I47V_I50V_I54L_L90M | -2.558541675 |
| I47V_I50V_L90M | -2.559058037 |
| V32I_M46I_I54M_I84V | -2.559873994 |
| M46I_I50V_L90M | -2.559943599 |
| V32I_I47V_I50V_I54L_L76V_V82T_I84V | -2.560124294 |
| V32I_I47V_I54M_V82T | -2.560232931 |
| M46I_I54L_L76V_V82F | -2.560729122 |
| M46I_I50V_I54M_L76V_V82F | -2.560730195 |
| V32I_T74P_L76V | -2.561029548 |
| V32I_I54L_L76V | -2.561754194 |
| V32I_I47V_I54M_V82T_I84V_L90M | -2.561958222 |
| M46I_I54L_L76V_V82T | -2.561992653 |
| V32I_I50V_I84V | -2.562062136 |

| | |
|---|---|
| M46I_I47V_I54L_L76V_V82F | -2.562731435 |
| I47V_I50V_I54L_V82F | -2.562908881 |
| M46I_I54M_T74P_I84V | -2.562920011 |
| M46I_I47V_I54M_I84V | -2.564810282 |
| V32I_M46I_I54M_L90M | -2.564862782 |
| M46I_I47V_I54M_L76V_V82F | -2.565337896 |
| V32I_M46I_I47V_I54L_L76V_I84V | -2.566624668 |
| M46I_I47V_I50V_I54L_I84V | -2.567569896 |
| V32I_M46I_I47V_V82F_I84V | -2.568816942 |
| V32I_I47V_L76V_V82F | -2.569088371 |
| M46I_I54L_L76V_V82T_I84V_L90M | -2.569458506 |
| V32I_M46I_I47V_I54L_V82T | -2.570851678 |
| V32I_M46I_I47V_I54L_L76V_V82T_I84V_L90M | -2.571494981 |
| M46I_I47V_I50V_I54M_V82F_I84V | -2.57156697 |
| M46I_I50V_I54M_L76V_V82T | -2.572623563 |
| I47V_L76V_V82F | -2.573695951 |
| V32I_M46I_I54L_L76V_I84V_L90M | -2.573695951 |
| V32I_M46I_I47V_I54M_L76V | -2.575096816 |
| M46I_I50V_I54L_L76V_V82T_I84V | -2.576375173 |
| M46I_I47V_I50V_L76V_V82T | -2.576673135 |
| V32I_M46I_I47V_V82F_I84V_L90M | -2.577202883 |
| I50V_V82T | -2.577478785 |
| V32I_M46I_I50V_I54M_L76V_V82F_I84V_L90M | -2.578203872 |
| I47V_I50V_I54M_L90M | -2.578334406 |
| V32I_M46I_I54M_L76V_V82F_I84V_L90M | -2.578388686 |
| V32I_V82F | -2.578540291 |
| I50V_I84V_L90M | -2.578647943 |
| V32I_M46I_I47V_I50V_I54M_V82F | -2.578985284 |
| I47V_I54M_V82F_I84V | -2.57956529 |
| V32I_I47V_I50V_I54M_V82F | -2.579648859 |
| M46I_I47V_V82F_I84V | -2.579687311 |
| V32I_I47V_I50V_I54L_V82T_I84V | -2.580929287 |
| V32I_I50V_I54M_I84V | -2.581432937 |
| V32I_M46I_I47V_I50V_V82T | -2.58144901 |
| M46I_I50V_L76V_V82F_I84V_L90M | -2.58150946 |
| I47V_I54M_L76V_I84V | -2.584939622 |
| V32I_M46I_I54M_L76V_I84V | -2.584939911 |
| V32I_M46I_V82T_L90M | -2.584939911 |
| I50V_I54M_L76V_V82F_I84V | -2.585551306 |
| V32I_M46I_I47V_I54M_L90M | -2.585565749 |
| V32I_M46I_I50V_I54M_L76V_I84V | -2.585637501 |
| V32I_I47V_I54M_I84V | -2.586015417 |
| I47V_I54M_L76V_V82F_L90M | -2.586855461 |

| | |
|---|---|
| V32I_I54L_V82F | -2.58728653 |
| V32I_I54M_V82T_I84V | -2.588110664 |
| V32I_M46I_I50V_I54M_L76V_V82T | -2.58924895 |
| V32I_I47V_I50V_I54M_I84V_L90M | -2.589716886 |
| I54L_L76V_I84V_L90M | -2.589966563 |
| M46I_I47V_I50V_I54L_L76V_I84V_L90M | -2.591698367 |
| V32I_I47V_I50V_L76V_L90M | -2.592783447 |
| M46I_I54M_L76V_L90M | -2.592949907 |
| M46I_I54L_I84V_L90M | -2.593273928 |
| I47V_I50V_L76V_V82F_I84V_L90M | -2.593487734 |
| I54M_L76V_L90M | -2.593667849 |
| V32I_M46I_I47V_I50V_L76V_V82F_I84V | -2.593770514 |
| V32I_M46I_I54M_L76V | -2.594195371 |
| V32I_L76V_V82T | -2.594308645 |
| V32I_M46I_I47V_I54M_V82T_I84V | -2.594323416 |
| I47V_L76V_I84V_L90M | -2.594488123 |
| V32I_I47V_I50V_I84V | -2.594511654 |
| M46I_I50V_I54L_L76V_V82F_L90M | -2.594775453 |
| M46I_I54M_L76V_V82F_I84V | -2.595742618 |
| V32I_I50V_I54L_L90M | -2.597472969 |
| V32I_I47V_I54M_L76V_V82T | -2.597949069 |
| V32I_I47V_I54L_V82F | -2.597992565 |
| V32I_M46I_I54M_V82F | -2.598509756 |
| V32I_I54M_L76V_V82F_I84V_L90M | -2.599097544 |
| V32I_M46I_I50V_I54L_V82T_I84V | -2.599848376 |
| M46I_I47V_I50V_V82F_I84V | -2.600472247 |
| V32I_M46I_I54L_L76V_I84V | -2.600502957 |
| M46I_I50V_V82T_L90M | -2.600667111 |
| M46I_I50V_L76V | -2.600932822 |
| I54M_I84V_L90M | -2.601304997 |
| V32I_M46I_I47V_I50V_L76V_V82F_L90M | -2.601458205 |
| I50V_I54M_L76V_I84V_L90M | -2.601838077 |
| V32I_I54L_I84V | -2.601991912 |
| V32I_I47V_I50V_I54L_I84V | -2.603350649 |
| V32I_M46I_I54M_L76V_V82F | -2.603354499 |
| V32I_M46I_I47V_I50V_I54L_L76V_V82F | -2.60384872 |
| I47V_I50V_I54L_I84V | -2.604192761 |
| V32I_M46I_I54L_V82T_I84V | -2.605085757 |
| M46I_I50V_L76V_I84V | -2.605369717 |
| M46I_I47V_I54L_V82F_I84V | -2.605998826 |
| V32I_I47V_I54M_L76V_V82T_L90M | -2.606920388 |
| V32I_I50V_I54L_V82F | -2.607117819 |
| V32I_M46I_I50V_L76V_V82F_L90M | -2.607643917 |

| | |
|---|---|
| V32I_M46I_I47V_I50V_L76V_V82F | -2.607725659 |
| V32I_L76V_V82T_I84V | -2.609018847 |
| I54L_L76V_I84V | -2.61008568 |
| V32I_M46I_I54L_L90M | -2.6103164 |
| I47V_I54L_L76V_V82F_I84V | -2.610681957 |
| M46I_I50V_I54L_I84V | -2.610884049 |
| M46I_I47V_I50V_I54L_L76V_L90M | -2.611003522 |
| I47V_L76V_I84V | -2.611027661 |
| M46I_I54M_V82F_L90M | -2.611632997 |
| I54L_L76V_V82T_L90M | -2.612022396 |
| V32I_I47V_I50V_L76V_V82F | -2.612410381 |
| V32I_M46I_I47V_I50V_I54M_L76V_V82F_I84V | -2.614325898 |
| I47V_I54M_I84V_L90M | -2.614346786 |
| V32I_I50V_I54M_L76V_V82F | -2.615389673 |
| I47V_I50V_I54M_L76V_V82T_I84V_L90M | -2.615764129 |
| I50V_I54M_L76V_V82F_I84V_L90M | -2.615764129 |
| I47V_I54M_T74P | -2.616357414 |
| M46I_L76V_V82T | -2.616539032 |
| V32I_I47V_I54M_L76V_L90M | -2.617127836 |
| I50V_I54L_V82T_L90M | -2.618464131 |
| V32I_I47V_I50V_I54M_L76V_L90M | -2.618975909 |
| V32I_I47V_I84V_L90M | -2.619231963 |
| M46I_I54L_V82T_I84V | -2.619655634 |
| V32I_I47V_I54L_V82T_I84V | -2.620097067 |
| I47V_I50V_I54M_I84V_L90M | -2.620343549 |
| M46I_I47V_L76V_V82F_I84V | -2.620589621 |
| M46I_I50V_I54L_V82T | -2.621374158 |
| V32I_M46I_T74P_V82F_I84V | -2.621945881 |
| I50V_I54L_L76V_L90M | -2.623039824 |
| I47V_I54L_T74P_V82F | -2.623202882 |
| M46I_I47V_L76V_V82T | -2.623377956 |
| V32I_M46I_I54M_L76V_V82F_I84V | -2.624002982 |
| V32I_I47V_I54L_L76V_V82F_I84V | -2.624086136 |
| V32I_V82F_I84V | -2.624969429 |
| I54M_L76V_I84V | -2.624993809 |
| M46I_I47V_I50V_L76V_V82T_L90M | -2.624993809 |
| M46I_I50V_I54L_L76V_V82T | -2.626931847 |
| M46I_I50V_I54M_V82F_L90M | -2.62695518 |
| I54M_V82F_I84V_L90M | -2.627834184 |
| V32I_M46I_L76V_V82F_I84V_L90M | -2.628545794 |
| I47V_I50V_L76V_I84V | -2.628575079 |
| V32I_M46I_I54M_V82T_L90M | -2.62857514 |
| V32I_I47V_I50V_I54L_L90M | -2.628840975 |

| | |
|---|---|
| M46I_I54M_L76V_V82T_I84V_L90M | -2.62949394 |
| V32I_M46I_I54L_V82T_L90M | -2.62967277 |
| I50V_I54M_V82T_L90M | -2.630004568 |
| V32I_I47V_I50V_V82T_L90M | -2.630004568 |
| I47V_I50V_I54M_L76V_V82F_I84V | -2.630047759 |
| M46I_I47V_I50V_I54M_L76V_I84V_L90M | -2.630644114 |
| V32I_I54L_L90M | -2.630791626 |
| V32I_I47V_I50V_I54L_V82T | -2.631382519 |
| I47V_I50V_L76V_V82T_I84V | -2.632512072 |
| V32I_M46I_I50V_I54M_V82T | -2.634460432 |
| V32I_M46I_I47V_V82T_L90M | -2.634604046 |
| V32I_L76V_V82F | -2.635042302 |
| M46I_I47V_L76V_V82T_I84V | -2.635177001 |
| V32I_V82T | -2.636063803 |
| V32I_L76V | -2.6362339 |
| M46I_I47V_I54L_L76V_V82F_I84V_L90M | -2.63709181 |
| V32I_I50V_L76V_V82T_L90M | -2.638441771 |
| V32I_M46I_I47V_T74P_L76V_L90M | -2.638441771 |
| V32I_I50V_I54L_L76V_I84V | -2.638517767 |
| I50V_I54M_L76V_L90M | -2.640778706 |
| M46I_I54M_V82F_I84V | -2.640978456 |
| M46I_I47V_T74P_V82F | -2.642384662 |
| V32I_M46I_I47V_I54M_V82F_I84V_L90M | -2.642589502 |
| V32I_M46I_I50V_I54L_L76V_V82T_L90M | -2.643792852 |
| V32I_I47V_I50V_L76V_V82F_I84V_L90M | -2.643792852 |
| V32I_M46I_I54L_L76V_V82F | -2.644499131 |
| M46I_I47V_I50V_I54M_L76V_V82T | -2.644658476 |
| I47V_I54M_L76V_V82F_I84V_L90M | -2.64675829 |
| M46I_I47V_T74P_V82T | -2.64685722 |
| V32I_I50V_I54M_L76V_V82F_I84V | -2.648157105 |
| M46I_I47V_T74P_I84V | -2.64820179 |
| I47V_I54M_L76V_I84V_L90M | -2.648737295 |
| V32I_M46I_I54L_V82F_I84V_L90M | -2.649760003 |
| V32I_I50V_V82F | -2.649763308 |
| I47V_T74P_L76V_V82F | -2.649778768 |
| I50V_I54M_V82T_I84V_L90M | -2.649981695 |
| M46I_I54L_L76V_V82F_I84V | -2.650820475 |
| I50V_I54M_L76V_V82T | -2.651131791 |
| I50V_L76V_V82T | -2.651276993 |
| I47V_I50V_I54L_L76V_V82F_I84V_L90M | -2.651537069 |
| M46I_I47V_I50V_I54M_L76V_L90M | -2.65205461 |
| V32I_M46I_I47V_I50V_V82T_L90M | -2.652839895 |
| V32I_M46I_I50V_L76V_V82T_L90M | -2.652974449 |

| | |
|---|---|
| V32I_M46I_I47V_L76V_V82T_I84V_L90M | -2.652995727 |
| V32I_I47V_I54L_L76V_V82F | -2.653076428 |
| V32I_M46I_I47V_I50V_V82F_I84V | -2.653130477 |
| V32I_I47V_I54M_L76V_V82T_I84V | -2.654542626 |
| V32I_I47V_I54L_L76V_V82T_I84V | -2.655130276 |
| M46I_I54M_V82F_I84V_L90M | -2.655636142 |
| V32I_I47V_I50V_I54M_L76V_V82T_I84V_L90M | -2.655815867 |
| V32I_I50V_I54M_L76V_L90M | -2.657156814 |
| V32I_I47V_I54M_L76V_I84V_L90M | -2.657156814 |
| I50V_I54M_L76V_V82F_L90M | -2.658477058 |
| V32I_V82T_I84V | -2.659069142 |
| V32I_M46I_I47V_I50V_I54L_V82F_L90M | -2.66020484 |
| V32I_I47V_I50V_V82F_I84V | -2.660866796 |
| V32I_I50V_I54L_L76V_V82F_I84V | -2.661181817 |
| V32I_M46I_I47V_I54L_L76V_V82F_I84V | -2.661990219 |
| M46I_I47V_I50V_L76V_V82F_L90M | -2.661990219 |
| V32I_M46I_I47V_I54M_L76V_V82F_I84V | -2.662094599 |
| V32I_M46I_I54M_L76V_I84V_L90M | -2.662780904 |
| M46I_I47V_I50V_I54L_L76V_V82T_L90M | -2.662885327 |
| V32I_I54L_L76V_V82F | -2.663502126 |
| V32I_I47V_I50V_I54L_I84V_L90M | -2.664249034 |
| V32I_M46I_I47V_I50V_I54L_I84V | -2.664615593 |
| V32I_I50V_I54M_V82F | -2.664646706 |
| M46I_L76V_V82T_I84V | -2.665353707 |
| M46I_I50V_I54L_L76V_V82T_I84V_L90M | -2.665891949 |
| M46I_I54L_T74P_L76V_V82F | -2.666693573 |
| V32I_I50V_I54L_V82T_L90M | -2.667239858 |
| M46I_I54M_V82T_L90M | -2.668579588 |
| V32I_I54L_L76V_V82T_I84V | -2.669362071 |
| I50V_I54L_L76V_V82F_I84V | -2.669723226 |
| V32I_M46I_I50V_I54L_L76V_L90M | -2.670121791 |
| M46I_I50V_I54L_L76V_I84V_L90M | -2.670121791 |
| V32I_T74P_V82F | -2.670867523 |
| M46I_I47V_I50V_I84V | -2.67104038 |
| I47V_I50V_I54L_V82T_I84V_L90M | -2.671682886 |
| M46I_L76V_V82T_L90M | -2.67252156 |
| I47V_I50V_I54M_V82T_L90M | -2.673081906 |
| M46I_I47V_L76V_V82F_I84V_L90M | -2.673161579 |
| I47V_I54L_L76V_V82T | -2.673454261 |
| V32I_M46I_I47V_I50V_I54L_L76V_V82F_I84V_L90M | -2.673527178 |
| V32I_I50V_I54M_V82F_I84V | -2.674956205 |
| V32I_M46I_I54M_V82T_I84V_L90M | -2.675025515 |
| V32I_M46I_I54M_I84V_L90M | -2.675818129 |

| | |
|---|---|
| V32I_M46I_I47V_I54M_L76V_V82F | -2.677013213 |
| V32I_M46I_V82F | -2.677585769 |
| V32I_M46I_I50V_I54M_L76V | -2.677624401 |
| M46I_I54L_T74P_V82F | -2.677733268 |
| M46I_I50V_L76V_I84V_L90M | -2.677733268 |
| I50V_L76V_I84V_L90M | -2.678200755 |
| V32I_I50V_V82F_I84V | -2.67861091 |
| I47V_I54M_V82F_L90M | -2.678896996 |
| I47V_I50V_I54L_V82F_I84V | -2.679131341 |
| V32I_M46I_I47V_L76V_V82T_I84V | -2.679647654 |
| I54M_V82F_L90M | -2.680483093 |
| I50V_V82T_I84V_L90M | -2.680639451 |
| L10F_V32I_I47V_T74P_V82F | -2.680639451 |
| I47V_I50V_L76V_V82F_I84V | -2.680897435 |
| V32I_I54L_L76V_I84V_L90M | -2.681215127 |
| I47V_I50V_I54M_L76V_V82F_L90M | -2.682710918 |
| M46I_I47V_I54M_V82T_I84V | -2.683709442 |
| V32I_M46I_I50V_I54L_V82F | -2.684415249 |
| V32I_M46I_I54M_L76V_V82T | -2.684936502 |
| V32I_M46I_I47V_I54L_V82F_I84V | -2.685985252 |
| V32I_I50V_L76V | -2.685986618 |
| M46I_I47V_V82F_I84V_L90M | -2.687307834 |
| I50V_I54L_V82F_I84V_L90M | -2.687723739 |
| I50V_I54M_L90M | -2.688034269 |
| V32I_I47V_I50V_I54L_L76V_I84V_L90M | -2.691113787 |
| V32I_M46I_I47V_T74P_V82F_I84V | -2.691238488 |
| V32I_M46I_I47V_I50V_I54M_I84V | -2.691776878 |
| M46I_T74P_V82F | -2.692269178 |
| V32I_M46I_I47V_L76V | -2.694063693 |
| V32I_M46I_I47V_I50V_I54M_V82F_I84V_L90M | -2.694945375 |
| M46I_I47V_I54L_I84V_L90M | -2.695011869 |
| M46I_I47V_I54M_L76V_V82T_I84V | -2.696191118 |
| V32I_M46I_I47V_I54M_L76V_I84V_L90M | -2.696433718 |
| M46I_I50V_I54L_V82F_I84V_L90M | -2.696905886 |
| I50V_V82T_I84V | -2.697015991 |
| V32I_M46I_I47V_I50V_I54L_V82F_I84V | -2.699250843 |
| V32I_M46I_I47V_I54M_V82F_L90M | -2.699250843 |
| I47V_I54M_L76V_V82T_I84V | -2.699844664 |
| M46I_I47V_I50V_I54L_V82T_L90M | -2.700799129 |
| V32I_M46I_I47V_I50V_I54L_L76V_V82T_I84V | -2.701104416 |
| V32I_M46I_I50V_I54L_V82T_I84V_L90M | -2.701222709 |
| I47V_I50V_I54L_L76V_I84V | -2.701394554 |
| V32I_I47V_I54L_L76V_L90M | -2.701622558 |

| | |
|---|---|
| I54M_V82T_L90M | -2.702677725 |
| V32I_I54M_V82F | -2.703284974 |
| I50V_L90M | -2.70411396 |
| V32I_M46I_I47V_L76V_I84V | -2.705156736 |
| V32I_M46I_I47V_I54M_L76V_V82T | -2.705713049 |
| M46I_I47V_I50V_L76V_V82F_I84V_L90M | -2.706844598 |
| I50V_L76V_V82F_L90M | -2.707628564 |
| V32I_M46I_I50V_L90M | -2.707628564 |
| I47V_I50V_I54M_L76V_L90M | -2.707913535 |
| M46I_I47V_I54M_T74P | -2.708549675 |
| M46I_I54M_V82T_I84V | -2.71009627 |
| V32I_I47V_I50V_V82F | -2.710872025 |
| V32I_I50V_I54M_V82T_I84V_L90M | -2.711673684 |
| V32I_M46I_I47V_V82F_L90M | -2.714037849 |
| V32I_I47V_I50V_I54M_L76V_V82F_L90M | -2.714037849 |
| V32I_M46I_I47V_I50V_V82F_I84V_L90M | -2.71426045 |
| V32I_I47V_I50V_I54M_L76V | -2.714655623 |
| V32I_M46I_I50V_I54M_V82T_L90M | -2.715148579 |
| M46I_I47V_I54L_V82T_I84V_L90M | -2.715148579 |
| V32I_M46I_I50V_I54M_V82F_L90M | -2.718426471 |
| V32I_M46I_I54L_I84V_L90M | -2.718426471 |
| V32I_M46I_I47V_I54L_L76V_V82T_I84V | -2.719692735 |
| I47V_I50V_I54L_L76V_V82F_I84V | -2.720839023 |
| V32I_M46I_I54L_L76V_V82F_I84V_L90M | -2.722633819 |
| V32I_I47V_I50V_I54M_L76V_I84V | -2.724314629 |
| V32I_I47V_L76V_V82T | -2.72492704 |
| I50V_I54L_L76V_V82F_I84V_L90M | -2.72632472 |
| M46I_I47V_I50V_I54L_V82T_I84V | -2.727122243 |
| V32I_M46I_I47V_I54L_L76V_V82T | -2.728023137 |
| I54L_V82T_L90M | -2.72868722 |
| V32I_M46I_I47V_I50V_I54M_V82T | -2.729319329 |
| M46I_I50V_I54M_L90M | -2.730001065 |
| V32I_I54L_L76V_V82F_L90M | -2.730482105 |
| V32I_I50V_L90M | -2.730635131 |
| I54L_I84V_L90M | -2.731050114 |
| V32I_M46I_I47V_I50V_L90M | -2.731550208 |
| V32I_M46I_I47V_L76V_V82T_L90M | -2.732572608 |
| V32I_M46I_I47V_I54M_V82F | -2.733304858 |
| V32I_I54L_V82T | -2.733621636 |
| M46I_I47V_I50V_I54M_I84V_L90M | -2.733821168 |
| I54L_L76V_V82F_L90M | -2.734834543 |
| V32I_I54M_L76V | -2.734972314 |
| M46I_I47V_I50V_I54L_I84V_L90M | -2.735172431 |

| | |
|---|---|
| V32I_I47V_L76V_V82F_L90M | -2.735776461 |
| M46I_I47V_I50V_I54M_L76V_I84V | -2.737985537 |
| V32I_I47V_I50V_L90M | -2.738115938 |
| V32I_I50V_L76V_I84V_L90M | -2.740670123 |
| M46I_I50V_L76V_L90M | -2.740670123 |
| V32I_I47V_I50V_I54M_L76V_V82F_I84V | -2.740845171 |
| V32I_I54L_L76V_L90M | -2.741020078 |
| M46I_I50V_I54L_L76V_V82F | -2.741323572 |
| M46I_I54M_L76V_I84V | -2.741585699 |
| V32I_I47V_I54M_V82T_L90M | -2.741734038 |
| V32I_I50V_I54L_L76V | -2.741797113 |
| V32I_M46I_I47V_I50V_L76V_V82T | -2.743367639 |
| V32I_I47V_I54L_L76V_V82T_I84V_L90M | -2.743367639 |
| V32I_M46I_I54M_V82F_I84V | -2.744052578 |
| I47V_L76V_V82F_I84V_L90M | -2.745561716 |
| I50V_I54L_I84V | -2.746061547 |
| I47V_I54L_V82F_L90M | -2.746669354 |
| V32I_M46I_I50V_V82F | -2.747051945 |
| V32I_M46I_I47V_I54M_L76V_L90M | -2.74728322 |
| I50V_I54L_L76V_V82T_I84V_L90M | -2.74758624 |
| V32I_I54L_L76V_V82T | -2.747772055 |
| V32I_M46I_I47V_I54L_L76V_V82T_L90M | -2.747967551 |
| I47V_I54M_L76V_V82T_L90M | -2.748377044 |
| V32I_I50V_V82F_I84V_L90M | -2.749577842 |
| I47V_I54L_I84V_L90M | -2.749845868 |
| M46I_I47V_T74P | -2.749998505 |
| I50V_L76V_V82F_I84V | -2.750276147 |
| M46I_I54L_T74P_L76V_I84V | -2.750276147 |
| V32I_I50V_I54M_V82T_L90M | -2.750680257 |
| V32I_M46I_I47V_I50V_I54L_V82F_I84V_L90M | -2.751019416 |
| V32I_M46I_I54L_V82T_I84V_L90M | -2.751223073 |
| I47V_L76V_V82F_L90M | -2.751355321 |
| V32I_M46I_I47V_I54L_V82T_L90M | -2.752905915 |
| I50V_I54M_V82F | -2.755252485 |
| M46I_V82T_I84V_L90M | -2.755403432 |
| I47V_T74P_V82F_I84V | -2.755599193 |
| V32I_M46I_I47V_I54L_V82T_I84V_L90M | -2.756810127 |
| M46I_I50V_V82F_L90M | -2.757430639 |
| V32I_V82T_L90M | -2.75931813 |
| V32I_M46I_T74P_L76V | -2.760352958 |
| M46I_I50V_L76V_V82T_L90M | -2.761892164 |
| I47V_I54L_L76V_L90M | -2.763015739 |
| V32I_M46I_I54L_V82F_L90M | -2.763951268 |

| | |
|---|---|
| I47V_I54L_V82T_I84V_L90M | -2.76425258 |
| V32I_I50V_L76V_V82F | -2.765055197 |
| I47V_V82T_L90M | -2.766181523 |
| V32I_M46I_I50V_I54L_L76V_V82F_I84V_L90M | -2.767488116 |
| V32I_I47V_I54M_V82F_I84V | -2.77036278 |
| V32I_I54M_I84V | -2.77082035 |
| V32I_I50V_I54M_V82F_I84V_L90M | -2.771019043 |
| V32I_M46I_I47V_I50V_I84V | -2.771163291 |
| I47V_I54L_V82T | -2.771174343 |
| V32I_M46I_I50V_I54M_V82F_I84V | -2.772706255 |
| V32I_M46I_I84V_L90M | -2.772972176 |
| V32I_M46I_I54M_V82F_I84V_L90M | -2.774320493 |
| M46I_I47V_I50V_V82F_I84V_L90M | -2.775166798 |
| I47V_I50V_I54L_L76V_V82T_I84V_L90M | -2.775166798 |
| V32I_I54M_V82T_L90M | -2.775919179 |
| V32I_I50V_I54L_V82T | -2.776623371 |
| V32I_I54L_I84V_L90M | -2.77769548 |
| V32I_I54L_V82F_I84V_L90M | -2.777710253 |
| I47V_V82F_L90M | -2.778643649 |
| M46I_I47V_L76V_V82F_L90M | -2.779375242 |
| V32I_I50V_L76V_V82T | -2.78194754 |
| I47V_I50V_L76V_V82T_I84V_L90M | -2.782095551 |
| M46I_I54L_V82F_L90M | -2.782104783 |
| M46I_I50V_I54L_I84V_L90M | -2.78253581 |
| V32I_I54M_V82F_I84V_L90M | -2.782583198 |
| V32I_I50V_I54L_V82F_I84V | -2.783255836 |
| I54M_L76V_V82F_I84V_L90M | -2.783793986 |
| V32I_M46I_I47V_I54M_L76V_V82T_L90M | -2.783878744 |
| I54M_L76V_I84V_L90M | -2.785702411 |
| M46I_I47V_I50V_I84V_L90M | -2.786285527 |
| V32I_I50V_I54M_L90M | -2.788312074 |
| V32I_M46I_I47V_I54M_V82F_I84V | -2.788610933 |
| I50V_V82F_I84V | -2.789033107 |
| V32I_I47V_I50V_I54L_V82F_I84V_L90M | -2.789638204 |
| V32I_I50V_I54L_V82F_I84V_L90M | -2.790230495 |
| V32I_I47V_I50V_I54M_L76V_V82T_L90M | -2.790521579 |
| I50V_L76V_L90M | -2.79154817 |
| V32I_I84V_L90M | -2.792120443 |
| V32I_I47V_I54L_L76V_V82T_L90M | -2.792587685 |
| M46I_I50V_I54M_L76V_V82F_I84V | -2.792640917 |
| I47V_I50V_I54L_L76V_V82F_L90M | -2.793343731 |
| M46I_I47V_V82F_L90M | -2.794284975 |
| M46I_I47V_I54L_L76V_I84V | -2.794609897 |

| | |
|---|---|
| I47V_I50V_V82F_L90M | -2.794976274 |
| M46I_I47V_L76V_V82T_I84V_L90M | -2.795471195 |
| V32I_M46I_I47V_I50V_I54L_L76V_V82F_I84V | -2.795581499 |
| I47V_I54L_V82T_I84V | -2.795995558 |
| I47V_I50V_L76V_V82F | -2.796043285 |
| I50V_I54L_V82F_L90M | -2.796188852 |
| V32I_I47V_I54M_L76V_V82F_I84V | -2.797094203 |
| V32I_M46I_I47V_L76V_V82F | -2.798720006 |
| M46I_I54L_V82F_I84V_L90M | -2.799494486 |
| V32I_I47V_I50V_I84V_L90M | -2.799494486 |
| V32I_M46I_I50V_I54L_I84V_L90M | -2.79950404 |
| V32I_M46I_I50V_I54L_V82F_I84V | -2.800458781 |
| V32I_I54M_L76V_I84V_L90M | -2.801400706 |
| I47V_I54L_L76V_I84V_L90M | -2.804340107 |
| M46I_I47V_I54M_L76V_V82T_I84V_L90M | -2.804611728 |
| I47V_I54M_V82T_L90M | -2.805578187 |
| V32I_M46I_V82T_I84V | -2.807350968 |
| I47V_L76V_V82T_I84V | -2.807538239 |
| V32I_I47V_L76V_V82F_I84V | -2.810302578 |
| I54M_L76V_V82F_L90M | -2.811786619 |
| V32I_M46I_L76V_V82T_L90M | -2.812038896 |
| M46I_I50V_I54L_V82T_L90M | -2.812038896 |
| M46I_I54L_L76V_V82T_I84V | -2.8152313 |
| V32I_I47V_I50V_L76V_I84V | -2.815912165 |
| V32I_I54M_I84V_L90M | -2.816933479 |
| M46I_I50V_L76V_V82F | -2.817002137 |
| L10F_I47V_T74P | -2.817208499 |
| M46I_I47V_I50V_I54L_V82T_I84V_L90M | -2.817638991 |
| M46I_I47V_I50V_L76V_V82T_I84V | -2.817700411 |
| M46I_I47V_I54L_V82T_L90M | -2.818044037 |
| V32I_I54M_V82F_I84V | -2.818605544 |
| M46I_I54M_L76V_V82F_L90M | -2.820626208 |
| V32I_M46I_I47V_T74P_V82F | -2.821198026 |
| M46I_I47V_I54M_I84V_L90M | -2.821542879 |
| M46I_I47V_I50V_I54L_V82F_I84V_L90M | -2.821852771 |
| V32I_I54M_L90M | -2.822798568 |
| V32I_M46I_I47V_I54L_V82F_L90M | -2.825184825 |
| V32I_I54M_L76V_V82F | -2.825723103 |
| V32I_M46I_I47V_T74P_L76V | -2.828838954 |
| V32I_M46I_I47V_I54L_L76V_V82F_L90M | -2.830165915 |
| V32I_M46I_I47V_I50V_I54L_L76V_I84V | -2.831864409 |
| V32I_I50V_V82F_L90M | -2.833050173 |
| V32I_I54L_L76V_V82F_I84V | -2.833052959 |

| | |
|---|---|
| M46I_I47V_L76V_L90M | -2.833586822 |
| V32I_I50V_I84V_L90M | -2.833963757 |
| V32I_M46I_I47V_I54M_L76V_V82F_I84V_L90M | -2.834386778 |
| L10F_V32I_T74P | -2.834442082 |
| V32I_M46I_I47V_I54M_I84V_L90M | -2.834736416 |
| V32I_M46I_I50V_V82F_I84V | -2.836078327 |
| M46I_I50V_I54M_V82T_L90M | -2.837612878 |
| V32I_I47V_I50V_V82T | -2.841260915 |
| M46I_I54M_L76V_V82T_L90M | -2.841811338 |
| V32I_I47V_I54L_T74P_V82F_I84V | -2.843099868 |
| V32I_I47V_I50V_I54L_L76V_V82F_L90M | -2.844338519 |
| I47V_I50V_I54L_V82T_L90M | -2.84621305 |
| V32I_I47V_I50V_I54L_L76V_V82F_I84V_L90M | -2.846634965 |
| I47V_I54L_L76V_V82T_I84V_L90M | -2.847731928 |
| M46I_I47V_I50V_V82F_L90M | -2.848774364 |
| M46I_L76V_I84V_L90M | -2.85204933 |
| I47V_I54L_L76V_V82T_L90M | -2.853009306 |
| I47V_I50V_I54L_L76V_V82F | -2.854258065 |
| V32I_M46I_I47V_I54M_V82T_I84V_L90M | -2.854292014 |
| M46I_I47V_I50V_L90M | -2.85448195 |
| I50V_I54L_V82T_I84V_L90M | -2.855327282 |
| V32I_M46I_I47V_I50V_I54L_I84V_L90M | -2.85673071 |
| M46I_I54L_L76V_V82F_I84V_L90M | -2.85673071 |
| M46I_I50V_V82T_I84V_L90M | -2.85673071 |
| V32I_M46I_I50V_I54M_L76V_V82F_L90M | -2.85673071 |
| V32I_M46I_I47V_I54L_L76V_L90M | -2.856819709 |
| V32I_I50V_I54L_I84V_L90M | -2.857894065 |
| M46I_I47V_I54L_L76V_L90M | -2.860024969 |
| M46I_I47V_I54M_V82F_I84V_L90M | -2.861859827 |
| I47V_V82T_I84V_L90M | -2.863434375 |
| I50V_L76V_I84V | -2.864722617 |
| V32I_M46I_I50V_L76V_V82T | -2.865712708 |
| V32I_I50V_I54L_V82T_I84V | -2.866305969 |
| V32I_M46I_I47V_L76V_L90M | -2.866886362 |
| M46I_I47V_I54L_L76V_I84V_L90M | -2.869753618 |
| V32I_M46I_I54L_L76V_V82F_L90M | -2.871036634 |
| I47V_I54M_V82F_I84V_L90M | -2.871565275 |
| V32I_I47V_I54L_L76V_I84V | -2.874292049 |
| V32I_I47V_I50V_I54L_L76V_V82F_I84V | -2.874605418 |
| V32I_M46I_I54L_L76V_L90M | -2.876196794 |
| I47V_I50V_I54L_V82T_I84V | -2.878744795 |
| M46I_I54M_L76V_I84V_L90M | -2.881819735 |
| I47V_I50V_I84V_L90M | -2.881882498 |

| | |
|---|---|
| V32I_M46I_I50V_I54L_I84V | -2.882841838 |
| L10F_I47V_V82F | -2.885759791 |
| M46I_I50V_I54L_V82F | -2.886137456 |
| L10F_V32I_I47V_V82F | -2.887573186 |
| M46I_I47V_I54M_V82T_L90M | -2.888366517 |
| I47V_I50V_I54L_L76V_V82T_L90M | -2.888763788 |
| V32I_I50V_V82T | -2.889830248 |
| M46I_I50V_I54L_L76V_V82F_I84V | -2.891840254 |
| I47V_L76V_L90M | -2.892050675 |
| M46I_I47V_L76V_V82T_L90M | -2.89413112 |
| I47V_I50V_L76V_V82T_L90M | -2.89451773 |
| V32I_I47V_I50V_I54M_V82T_I84V | -2.897451109 |
| I54L_L76V_V82T_I84V_L90M | -2.898456637 |
| M46I_L76V_L90M | -2.899474971 |
| I47V_I50V_I54M_V82F_I84V_L90M | -2.901391673 |
| M46I_I50V_I54L_V82F_L90M | -2.902528985 |
| I50V_I54L_L76V_V82T_L90M | -2.909494886 |
| V32I_I47V_I50V_I54M_V82F_I84V_L90M | -2.909494886 |
| M46I_I50V_I84V_L90M | -2.912033518 |
| I47V_I50V_I54L_L76V_L90M | -2.91350245 |
| I47V_I54M_L76V_L90M | -2.913588074 |
| I47V_I50V_L76V_V82F_L90M | -2.914440045 |
| V32I_I54L_L76V_V82T_I84V_L90M | -2.914440045 |
| M46I_I47V_I50V_I54L_L76V_V82F_L90M | -2.915473365 |
| I50V_V82F_L90M | -2.91618681 |
| V32I_I47V_I50V_V82F_I84V_L90M | -2.916794124 |
| I47V_I50V_I54L_L90M | -2.919795708 |
| V32I_M46I_I47V_I54L_I84V_L90M | -2.920159203 |
| V32I_M46I_I47V_I54M_V82T_L90M | -2.920275247 |
| V32I_M46I_I50V_I54L_L76V_I84V | -2.921306311 |
| V32I_I47V_I54M_L76V_V82F_L90M | -2.924220277 |
| V32I_M46I_I54M_L76V_V82T_L90M | -2.925969296 |
| V32I_V82T_I84V_L90M | -2.926315803 |
| M46I_I47V_I50V_I54L_V82F_L90M | -2.926650706 |
| I50V_I54L_L76V_V82F_L90M | -2.928300485 |
| V32I_I50V_V82T_I84V | -2.929348222 |
| I47V_I50V_I54L_V82F_I84V_L90M | -2.930314889 |
| V32I_M46I_L76V_L90M | -2.930512184 |
| V32I_M46I_I47V_I84V_L90M | -2.930767442 |
| V32I_L76V_V82F_I84V_L90M | -2.931322301 |
| V32I_I54M_V82T_I84V_L90M | -2.93168591 |
| V32I_I50V_I54L_L76V_V82F | -2.932172194 |
| L10F_T74P | -2.93257316 |

| | |
|---|---|
| I50V_V82T_L90M | -2.933296637 |
| M46I_I47V_I50V_I54M_L76V_V82F | -2.935110164 |
| M46I_I54L_L76V_L90M | -2.936265004 |
| V32I_L76V_I84V_L90M | -2.937386958 |
| V32I_I50V_V82T_L90M | -2.944121959 |
| L10F_V32I_V82F | -2.944948233 |
| V32I_I47V_I54L_L76V_V82F_I84V_L90M | -2.949210554 |
| V32I_I47V_I50V_L76V_V82F_I84V | -2.949799739 |
| V32I_M46I_I54L_L76V_V82T_I84V_L90M | -2.949799739 |
| L10F_V82F | -2.952585865 |
| V32I_I47V_I50V_I54M_L76V_V82T | -2.952672702 |
| V32I_V82F_I84V_L90M | -2.953355249 |
| V32I_I47V_I54L_L76V_V82F_L90M | -2.956161912 |
| V32I_M46I_I50V_V82F_I84V_L90M | -2.956845863 |
| I47V_I54L_L76V_V82F_L90M | -2.957205686 |
| I47V_I54L_V82F_I84V_L90M | -2.958304273 |
| V32I_M46I_I47V_I50V_L76V_V82F_I84V_L90M | -2.958304273 |
| V32I_I47V_L76V_V82T_I84V_L90M | -2.961082916 |
| V32I_I47V_V82F_I84V_L90M | -2.961705783 |
| V32I_I50V_I54M_I84V_L90M | -2.961705783 |
| V32I_I47V_I54L_V82T_I84V_L90M | -2.964403333 |
| V32I_M46I_I47V_I50V_I54L_V82T_I84V_L90M | -2.96527903 |
| I47V_I50V_V82T_L90M | -2.965812221 |
| V32I_M46I_L76V_V82T | -2.967198724 |
| V32I_I50V_I54L_L76V_L90M | -2.967538841 |
| V32I_M46I_I54M_V82F_L90M | -2.96827681 |
| I47V_I54M_L76V_V82T_I84V_L90M | -2.968460911 |
| M46I_L76V_V82F_L90M | -2.968842107 |
| V32I_M46I_I47V_L76V_V82F_I84V | -2.969358578 |
| V32I_I47V_I50V_I54L_V82T_I84V_L90M | -2.96943499 |
| V32I_I47V_I50V_L76V_V82F_L90M | -2.971523553 |
| V32I_M46I_I50V_I54L_L76V_V82T_I84V | -2.973613286 |
| V32I_L76V_V82F_I84V | -2.974307644 |
| M46I_I47V_I54M_V82F_L90M | -2.974997983 |
| V32I_L76V_V82T_L90M | -2.975581131 |
| I47V_I50V_I54L_V82F_L90M | -2.975643652 |
| V32I_I54L_L76V_V82F_I84V_L90M | -2.976143881 |
| V32I_I47V_I54M_L76V_V82T_I84V_L90M | -2.977092043 |
| M46I_I54L_L76V_V82F_L90M | -2.977491965 |
| I50V_L76V_V82F_I84V_L90M | -2.977629468 |
| V32I_M46I_I47V_I54L_L76V_V82F_I84V_L90M | -2.983116582 |
| V32I_I54M_V82F_L90M | -2.983740914 |
| V32I_M46I_I47V_I54L_V82F_I84V_L90M | -2.986466275 |

| | |
|---|---|
| V32I_M46I_I54M_L76V_L90M | -2.989638376 |
| V32I_I47V_L76V_V82T_L90M | -2.991505971 |
| M46I_I47V_I54L_V82F_L90M | -2.992326002 |
| V32I_I47V_I54L_V82F_L90M | -2.994754109 |
| V32I_M46I_I50V_I54L_V82F_L90M | -2.995070691 |
| V32I_I47V_I50V_I54M_L76V_V82F_I84V_L90M | -2.995070691 |
| M46I_I47V_V82T_I84V_L90M | -3.002118488 |
| V32I_I54L_V82F_L90M | -3.0073194 |
| V32I_I47V_I54L_I84V_L90M | -3.008063425 |
| L10F_V32I_T74P_V82F | -3.00843883 |
| V32I_I54M_L76V_V82F_L90M | -3.009884011 |
| I47V_I50V_L76V_L90M | -3.011231501 |
| V32I_M46I_I50V_I54L_L90M | -3.01270368 |
| V32I_M46I_I47V_I50V_I54L_L76V_V82T_I84V_L90M | -3.01270368 |
| I47V_I50V_V82T_I84V_L90M | -3.015784299 |
| V32I_I47V_I54M_V82F_I84V_L90M | -3.019456466 |
| I50V_V82F_I84V_L90M | -3.019624045 |
| V32I_M46I_I47V_I50V_L76V_L90M | -3.020127698 |
| V32I_I47V_V82T_L90M | -3.022150318 |
| I47V_I54L_V82T_L90M | -3.02577053 |
| I54L_V82F_I84V_L90M | -3.028088441 |
| M46I_I47V_I54M_L76V_I84V_L90M | -3.034605521 |
| V32I_M46I_I47V_L76V_V82F_I84V_L90M | -3.034605521 |
| I47V_I54L_L76V_V82F_I84V_L90M | -3.035229313 |
| V32I_I47V_I54M_L76V_V82F_I84V_L90M | -3.037456063 |
| V32I_I47V_I54M_I84V_L90M | -3.039941832 |
| I47V_I50V_L76V_I84V_L90M | -3.040140351 |
| V32I_I47V_V82F_L90M | -3.040890863 |
| V32I_I47V_I50V_I54L_V82T_L90M | -3.041667376 |
| V32I_I47V_I54L_L76V_V82T | -3.046813137 |
| I50V_L76V_V82T_L90M | -3.048616236 |
| M46I_I47V_I54M_L76V_V82F_L90M | -3.059672932 |
| I47V_V82F_I84V_L90M | -3.061066858 |
| V32I_M46I_V82F_L90M | -3.064048986 |
| V32I_I54L_L76V_I84V | -3.067381112 |
| V32I_I54M_L76V_L90M | -3.075350489 |
| M46I_I54M_L76V_V82F_I84V_L90M | -3.083001887 |
| M46I_I50V_I54M_L76V_L90M | -3.083001887 |
| M46I_I47V_I54L_V82F_I84V_L90M | -3.085545277 |
| V32I_I54M_L76V_V82T_L90M | -3.092568227 |
| I47V_I54M_V82T_I84V_L90M | -3.092662305 |
| V32I_I54L_V82T_I84V_L90M | -3.092939102 |
| V32I_I50V_L76V_L90M | -3.094373726 |

| | |
|---|---|
| V32I_M46I_I47V_I50V_V82F_L90M | -3.097991882 |
| V32I_M46I_I54L_L76V_V82T_L90M | -3.106988055 |
| V32I_I47V_I54L_V82T_L90M | -3.110262449 |
| V32I_I47V_L76V_V82F_I84V_L90M | -3.115532432 |
| I47V_I50V_V82F_I84V_L90M | -3.117467829 |
| V32I_I47V_I50V_V82F_L90M | -3.118730406 |
| V32I_I47V_L76V_I84V_L90M | -3.122228022 |
| M46I_I47V_I54L_L76V_V82F_L90M | -3.126066445 |
| I47V_L76V_V82T_L90M | -3.12986895 |
| V32I_I47V_L76V_L90M | -3.138057181 |
| I54L_L76V_V82F_I84V_L90M | -3.176132395 |
| V32I_M46I_I47V_L76V_V82F_L90M | -3.181241801 |
| V32I_L76V_L90M | -3.182357104 |
| V32I_V82F_L90M | -3.232583435 |
| V32I_I47V_I54L_V82F_I84V_L90M | -3.243090089 |
| V32I_I54L_V82T_L90M | -3.265504336 |
| L10F_T74P_V82F | -3.338735946 |
