## Supplementary file 2 for "Predominance of positive epistasis among drug resistance-associated mutations in HIV-1 protease"

Supplementary Table 2: Sequence of oligonucleotides used in this research.

| oligonucleotides | sequence |
|---|---|
| O1F | ACTCTTTGGCAGCGACCC**YTC**GTCACAATAAAGATAGGGGGGCAATTAAAGGAAGCT |
| O1R | /5Phos/TCCTTTAATTGCCCCCCTATCTTTATTGTGAC**GAR**GGGTCGCTGCCAAAGAGT |
| O2F | /5Phos/CTATTAGATACAGGAGCAGATGATACA**RTA**TTAGAAGAAATGAATTTGCCAGGA |
| O2R | TCCTGGCAAATTCATTTCTTCTAA**TAY**TGTATCATCTGCTCCTGTATCTAATAGAGCT |
| O3F1 | TGGAAACCAAAA**ATSRTA**GGGGGA**RTT**GGAGGTTTT**ATC**AAAGTAAGACAGTATGAT |
| O3F2 | TGGAAACCAAAA**ATSRTA**GGGGGA**RTT**GGAGGTTTT**ATG**AAAGTAAGACAGTATGAT |
| O3F3 | TGGAAACCAAAA**ATSRTA**GGGGGA**RTT**GGAGGTTTT**CTC**AAAGTAAGACAGTATGAT |
| O3R1 | /5Phos/TACTGTCTTACTTT**GAT**AAAACCTCC**AAY**TCCCCC**TAYSAT**TTTTGGTTTCCA |
| O3R2 | /5Phos/TACTGTCTTACTTT**GAT**AAAACCTCC**AAY**TCCCCC**TAYSAT**TTTTGGTTTCCA |
| O3R3 | /5Phos/TACTGTCTTACTTT**GAT**AAAACCTCC**AAY**TCCCCC**TAYSAT**TTTTGGTTTCCA |
| O4F | /5Phos/CAGATACTCATAGAAATCTGCGGACATAAAGCTATAGGT**MCA**GTA**KTA**GTAGGA |
| O4R | TCCTAC**TAM**TAC**TGK**ACCTATAGCTTTATGTCCGCAGATTTCTATGAGTATCTGATCA |
| O5F1 | /5Phos/GACCTACACCT**GTC**AAC**RTA**ATTGGAAGAAATCTG**WTG**ACTCAGATTGGCTGCACTTTA |
| O5F2 | /5Phos/GACCTACACCT**ACC**AAC**RTA**ATTGGAAGAAATCTG**WTG**ACTCAGATTGGCTGCACTTTA |
| O5F3 | /5Phos/GACCTACACCT**TTC**AAC**RTA**ATTGGAAGAAATCTG**WTG**ACTCAGATTGGCTGCACTTTA |
| O5R1 | TAAAGTGCAGCCAATCTGAGT**CAW**CAGATTTCTTCCAAT**TAY**GTT**GAC**AGGTGTA |
| O5R2 | TAAAGTGCAGCCAATCTGAGT**CAW**CAGATTTCTTCCAAT**TAY**GTT**GAC**AGGTGTA |
| O5R3 | TAAAGTGCAGCCAATCTGAGT**CAW**CAGATTTCTTCCAAT**TAY**GTT**GAC**AGGTGTA |
| Recover_O1+O2_F | ACTCTTTGGCAGCGACCC |
| Recover_O1+O2_R | CCGGTCTCTCTTCCTGGCAAATTCATTTCTTC |
| Recover_O3+O4_F | TGGAAACCAAAAATSRTAGGG |
| Recover_O3+O4_R | CCGGTCTCAGGTCCTACTAMTACTGKACCTATAGC |
| Recover_O345_F | CCGGTCTCGGAAGATGGAAACCAAAAATSRTAGGG |
| Recover_O345_R | TAAAGTGCAGCCAATCTGAGT |
| Up_ApaI_F | GCAGGGCCCCTAGGAAA |
| Up_R | GGGTCGCTGCCAAAGAGT |
| Down_F | ACTCAGATTGGCTGCACTTTA |
| Down_SbfI_R | AACCCTGCAGGATGTGG |
| Mut1_L90M_F | GAAGAAATCTGATGACTCAGATTGG |
| Mut2_V82T_F | CACCTACCAACATAATTGGAAGAAATC |
| Mut3_V82F_F | CACCTTTCAACATAATTGGAAGAAATC |
| Mut4_L76V_F | CTATAGGTACAGTAGTAGTAGGACCTAC |
| Mut5_T74P_F | CTATAGGTCCAGTATTAGTAGGACCTAC |
| Mut6_I47V_F | GGAAACCAAAAATGGTAGGGGGAATTG |
| Mut7_L10F_F | CGACCCTTCGTCACAATAAAG |
| Mut1_L90M_R | CCAATCTGAGTCATCAGATTTCTTC |
| Mut2_V82T_R | GATTTCTTCCAATTATGTTGGTAGGTG |
| Mut3_V82F_R | GATTTCTTCCAATTATGTTGAAAGGTG |
| Mut4_L76V_R | GTAGGTCCTACTACTACTGTACCTATAG |
| Mut5_T74P_R | GTAGGTCCTACTAATACTGGACCTATAG |
| Mut6_I47V_R | CAATTCCCCCTACCATTTTTGGTTTCC |
| Mut7_L10F_R | CTTTATTGTGACGAAGGGTCG |
